# Heterogeneous Mediation Analysis on Epigenomic PTSD and Traumatic Stress in a Predominantly African American Cohort

**DOI:** 10.1101/2020.10.13.336826

**Authors:** Fei Xue, Xiwei Tang, Grace Kim, Karestan C. Koenen, Chantel L. Martin, Sandro Galea, Derek Wildman, Monica Uddin, Annie Qu

**Affiliations:** Purdue University; University of Virginia; University of Illinois College of Medicine; Harvard T.H. Chan School of Public Health; The University of North Carolina at Chapel Hill; Boston University; University of South Florida; University of California Irvine

**Keywords:** Clustering, difference of convex, DNA methylation, high-dimensional mediators, linear structural equation modeling, variable selection

## Abstract

DNA methylation (DNAm) has been suggested to play a critical role in post-traumatic stress disorder (PTSD), through mediating the relationship between trauma and PTSD. However, this underlying mechanism of PTSD for African Americans still remains unknown. To fill this gap, in this paper, we investigate how DNAm mediates the effects of traumatic experiences on PTSD symptoms in the Detroit Neighborhood Health Study (DNHS) (2008–2013) which involves primarily African Americans adults. To achieve this, we develop a new mediation analysis approach for high-dimensional potential DNAm mediators. A key novelty of our method is that we consider heterogeneity in mediation effects across sub-populations. Specifically, mediators in different sub-populations could have opposite effects on the outcome, and thus could be difficult to identify under a traditional homogeneous model framework. In contrast, the proposed method can estimate heterogeneous mediation effects and identifies sub-populations in which individuals share similar effects. Simulation studies demonstrate that the proposed method outperforms existing methods for both homogeneous and heterogeneous data. We also present our mediation analysis results of a dataset with 125 participants and more than 450, 000 CpG sites from the DNHS study. The proposed method finds three sub-groups of subjects and identifies DNAm mediators corresponding to genes such as *HSP90AA1* and *NFATC1* which have been linked to PTSD symptoms in literature. Our finding could be useful in future finer-grained investigation of PTSD mechanism and in the development of new treatments for PTSD.

## 1 Introduction

Post-traumatic stress disorder (PTSD) is a serious mental health disorder that people may develop after they experience or witness a traumatic event, such as a natural disaster, a serious accident, a war, or sexual violence. People suffering from PTSD have symptoms such as disturbing thoughts, nightmares related to the events, mental or physical distress to trauma-related cues, attempts to avoid trauma-related cues, and negative alterations in thinking and feeling (Morrison et al., 2019). As shown in Roberts et al. (2011) and Himle et al. (2009), the prevalence of PTSD is higher in African Americans (AAs) than whites. This is possibly due to that PTSD is an adversity-related mental disorder and that AAs are more likely to encounter socially adverse experiences of discrimination and isolation which have a profound impact on mental health (Hudson et al., 2013; Cacioppo et al., 2015). However, research for the course of PTSD among AAs is very limited.

As suggested by Rusiecki et al. (2013) and Morrison et al. (2019), DNA methylation (DNAm), an epigenetic mechanism associated with the regulation of gene expression, plays a critical role in the pathophysiology of the PTSD. Since DNAm is a reversible process (Ramchandani et al., 1999), uncovering the role of DNAm in the pathophysiology of PTSD may facilitate the development of new potential treatments for PTSD (Rusiecki et al., 2013). For example, the epigenetics literature suggests that DNAm could mediate the effect of traumatic events on depression (Gao et al., 2019; Vangeel et al., 2015; Dempster et al., 2014; Tyrka et al., 2016; Januar et al., 2015; Schuster et al., 2017; Zhao et al., 2013; Lei et al., 2015; Turecki and Meaney, 2016; van der Knaap et al., 2015). In particular, Peng et al. (2018) discover that DNAm levels at two cytosine-phosphate-guanine (CpG) probes mediate the effects from childhood trauma to depressive symptoms. Rutten et al. (2018) also observe that DNAm mediates the relationship between combat trauma and PTSD symptoms.

Moreover, according to Dickstein et al. (2010), heterogeneity occurs in the course of PTSD. For example, Kim et al. (2019b) observe gender differences in risk of PTSD, indicating a potential mechanism that yields heterogeneity across subjects in the effects of trauma. This motivates us to develop a heterogeneous model. In addition, Heinzelmann and Gill (2013) suggest that epigenetics may play a key role in the heterogeneous responses to trauma and differential risk of PTSD. Furthermore, Orcutt et al. (2004) and Dickstein et al. (2010) reveal distinct trajectories of PTSD symptoms. This is crucial since finding all the prototypical patterns of adaptation to trauma could help us identify biomarkers and risk factors for PTSD. Nevertheless, to the best of our knowledge, heterogeneity in the mediation effects of the DNAm between traumatic events and PTSD symptoms has not been investigated so far.

In fact, heterogeneous mediation effects could arise frequently due to factors such as demographic and genetic characteristics, medical history, lifestyle, and unobserved attributes of subjects. Mediators in different sub-populations could vary or have different effects on the outcome. For example, as illustrated in Figure 1, high pressure on subjects could mediate the effects from the exposure of competitive environments to the performance of subjects. Yet, high pressure may have positive effects on performance of subjects in some sub-population but negative effects for subjects in another sub-population. This heterogeneity among subjects likely comes from stress tolerance and the ability to turn pressure into power, which may vary greatly across different groups of people. In this case, it could be infeasible to identify the true mediator in a homogeneous model, since opposite effects could be canceled out. Recently, several mediation methods have been studied for heterogeneous mediation effects (Qin and Hong, 2017; Dyachenko and Allenby, 2018). However, they either require pre-specified sub-populations or only focus on a single mediator variable.

**Figure 1:**
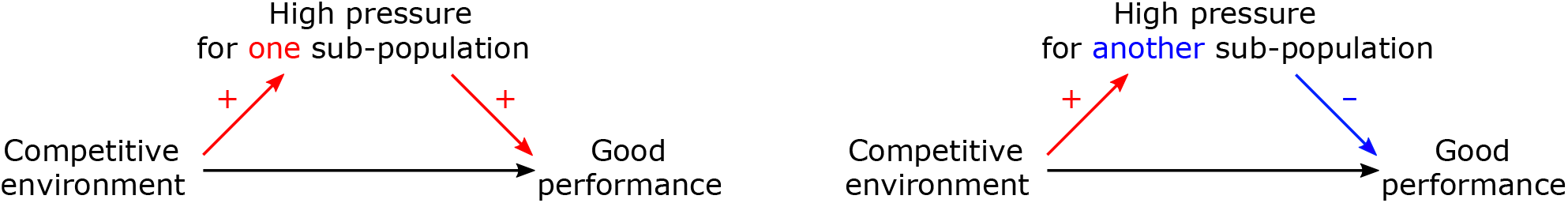
A heterogeneous mediation example.

In this paper, we aim to investigate the high-dimensional heterogeneous mediation effects of DNAm variation on the relationship between trauma exposures and PTSD symptoms using the Detroit Neighborhood Health Study (DNHS) data (Uddin et al., 2010). The DNHS is a representative longitudinal cohort study involving primarily African Americans. In addition, we propose a novel mediator selection method which can identify subgroups of subjects and select mediators in each subgroup simultaneously for high-dimensional potential mediators. We refer to this new method as “the proposed method” in the rest of this paper.

Note that the goal of our study is not to compare differences in PTSD risk by race and thus an interaction between race and trauma is not needed in our model. Rather, AAs’ higher prevalence of PTSD is a motivation for us to examine the mediation effects in a sample from this population. Hence the focus of this paper is to investigate the mediation mechanism in a predominantly AA sample (the DNHS data). On the other hand, in fact, there are few non-AA subjects in the DNHS dataset, indicating that it is inappropriate to use this dataset to compare racial differences in PTSD.

Our numerical studies show that the proposed method outperforms existing homogeneous methods in terms of mediation effect estimation and mediator selection. More importantly, the proposed method identifies meaningful DNAm mediators which are not selected by the homogeneous mediation methods on the DHNS data. Specifically, the selected DNAm CpG probes correspond to genes including *HSP90AA1, SMARCA4*, and *NFATC1* which are indeed associated with PTSD risk (Criado-Marrero et al., 2018; Breen et al., 2019; Kim et al., 2019a; Raabe and Spengler, 2013; Kim et al., 2019a; Kuan et al., 2017). These DNAm mediators can be highly informative in future development of novel interventions for PTSD. In addition, our data analysis shows the potential heterogeneity in mediation effects of DNAm on PTSD risk, which suggests finer-grained comparisons in future PTSD research.

The remainder of this paper is organized as follows. In Section 2, we describe details of the DNHS data which we analyze in this article. In Section 3, we propose the heterogeneous mediation method and illustrate the implementation of the proposed method. Section 4 provides numerical studies through simulations. In Section 5, we apply the proposed method to the DNHS dataset. Finally, we conclude this study with discussion in Section 6.

## 2 DNHS Data

Our work is motivated by the Detroit Neighborhood Health Study (DNHS) where samples are collected between 2008–2013 from predominantly African American (AA) adults living in Detroit, Michigan. Studies suggest that DNA methylation (DNAm) could play a crucial role as mediators in the underlying relationship between traumatic events and PTSD (Gao et al., 2019; Vangeel et al., 2015; Dempster et al., 2014; Tyrka et al., 2016). Our work was further inspired by initial joint analysis of expression and EWAS data in GRRN genes (Vukojevic et al., 2014; Palma-Gudiel et al., 2015; Kim et al., 2019a; Wani et al., 2021), to find potential functional significance, and the DNHS is one of the few available datasets with both of these data types (Uddin et al., 2010).

In this paper, in order to understand the underlying mechanism of PTSD in AAs, we conduct mediation analysis of DNAm on the relationship between trauma exposures and PTSD symptoms. One significant impact of identifying true mediators is that the occurrence of PTSD can be potentially intervened through the DNAm mediators. This is especially important for PTSD patients in the DNHS since the independent variables such as the previous trauma experience and adverse events cannot be altered after.

The DNHS is comprised of five survey waves where a total of 2081 participants have completed a 40-minute telephone survey. The survey includes questions on participants’ neighborhoods, mental and physical health status, social support, exposure to traumatic events, PTSD symptoms, and various demographics characteristics. All participants have been offered an opportunity to provide a blood specimen for genetic testing of DNA. In particular, the DNHS measures the PTSD symptom severity of participants through the widely-used self-report PTSD Checklist, Civilian Version (PCL-C) (Blanchard et al., 1996; Grubaugh et al., 2007). The PCL-C set contains 17 items corresponding to key symptoms of PTSD. Participants indicate how much they have been bothered by each symptom using a 5-point (1–5) scale in reference to their worst traumatic experience. To access the overall severity, we calculate the average of the 17 items and treat the average PCL-C score as a representative of the PTSD symptom severity for each participant.

The DNHS also records the types of traumatic or stressful events that each participant has experienced. In our survey, we have specific questions such as “Have you experienced exposure to a war zone in the military or as a civilian?” to understand the exact nature of trauma exposures. Thus, a trauma exposure of a subject provides information of a certain type of trauma. We calculate the total number of trauma exposures (i.e., how many different types of traumas a subject experienced) and use it as a trauma feature characterizing overall trauma severity for each subject. Here we use the total number of experienced event types since many studies have used it as a measure of severity of trauma, e.g., Lee and Park (2018); Irish et al. (2013); Harte et al. (2015); Farley et al. (2001); Kessler et al. (2017).

The blood specimens in the DNHS are processed according to Weckle et al. (2015), and the DNA Mini Kit (Qiagen, Germantown, MD) is used to extract genomic DNA from peripheral blood. The extracted DNA samples are bisulfite-converted using the Zymo EZ-96 DNA methylation kit (Zymo Research, Irvine, CA). The converted samples are then profiled through the Illumina Infinium 450 K DNA methylation array (Illumina, San Diego, CA) according to manufacturer protocols. Specifically, we assess the methylation levels of more than 450,000 CpG sites which cover 99% of reference sequence (RefSeq) genes. More detailed explanation of the processing procedures are provided in Kim et al. (2019b); Ward-Caviness et al. (2020); Wolf et al. (2018); Ratanatharathorn et al. (2017), and Uddin et al. (2018).

There are several existing studies on the DNHS data from various aspects. Uddin et al. (2010) find that Detroit residents have PTSD prevalence more than twice of the corresponding prevalence in the entire US, and that a person’s immune-related functions are related to genes with relatively lower levels of methylation. McClure et al. (2018) assess the association between environmental stressors and the Great Recession, while Horesh et al. (2015) and Kim et al. (2019b) reveal gender differences in PTSD. Moreover, Chang et al. (2012); Ratanatharathorn et al. (2017); Uddin et al. (2018) and Nevell et al. (2014) study genetic factors associated with the risk of PTSD in the DNHS. However, none of them have conducted mediation analysis for relationship between traumatic exposures and PTSD symptoms on the DNHS data.

In this paper, we utilize the baseline wave in DNHS for our mediation analysis. In total, there are 125 subjects in the DNHS who have available PTSD measurements, trauma type numbers, and DNAm data. This sample size is relatively small. In fact, many existing studies on PTSD-related mediation analysis have around one hundred participants (Kwon et al., 2021; Ruhlmann et al., 2019; Demir et al., 2020; Kearney et al., 2013; van der Vleugel et al., 2020; Kelly et al., 2019). A relatively small sample size could be a challenge for existing statistical methods, which motivates us to develop more powerful statistical methods to identify mediators. The demographic characteristics of these DNHS participants are summarized in Table 1. The study participants are predominantly (88.8%) African-American (AA). In addition, of the 125 participants, 64.8% are female, and 43.2% are current smoking, referring to any cigarette smoking in the past 30 days. The age of DNHS participants has a wide range from 20 to 89 years with a median of 53 years.

**Table 1:**
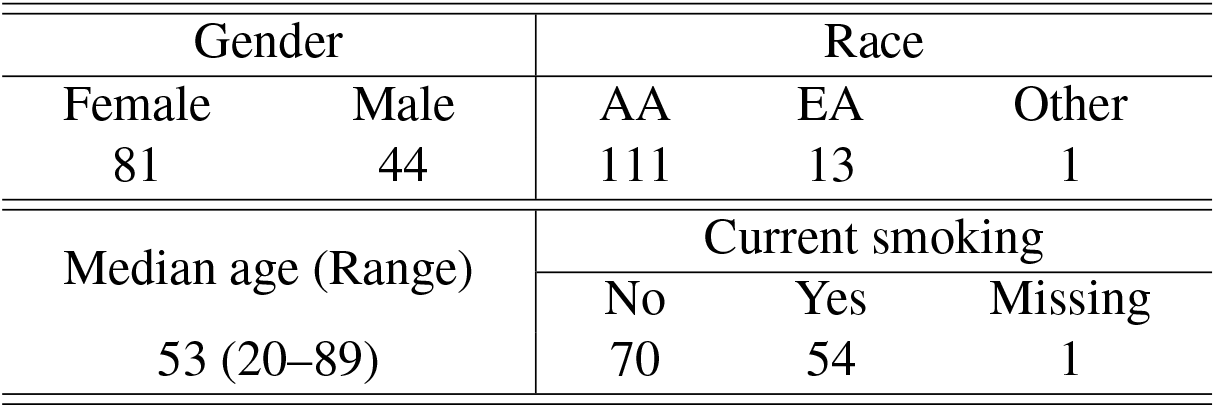
Key demographic characteristics. “AA” represents African American. “EA” represents European American. “Current smoking” refers to any cigarette smoking in the past 30 days.

| Gender |  | Race |  |  |
| --- | --- | --- | --- | --- |
| Female | Male | AA | EA | Other |
| 81 | 44 | 111 | 13 | 1 |
| Median age (Range) |  | Current smoking |  |  |
|  |  | No | Yes | Missing |
| 53 (20–89) |  | 70 | 54 | 1 |

Studies show that the pathophysiology of PTSD is associated with DNAm in glucocorticoid receptor regulatory network (GRRN) genes (Rusiecki et al., 2013). Therefore, we screen DNAm CpG probes via an expression quantitative trait methylation (eQTM) analysis on GRRN-annotated DNAm CpG probes and 53 expressed GRRN genes. Here these 53 genes correspond to 1, 680 GRRN-annotated probes. For each probe, we examine whether the probe is significantly correlated with expression levels of the corresponding gene at a significance level of 0.05. If it is significant, we select that probe; otherwise, the probe is not selected. That is, we choose the CpG probes in probe-gene pairs with significant p-values. Through this, we identify 144 CpG probes significantly correlated with the GRRN genes. For mediation analysis, we treat the PCL-C score as a dependent variable, the total number of trauma exposures as an independent variable, and the 144 DNAm CpG probes as potential mediators. Our aim is to select key mediators between the independent and dependent variables from the 144 DNAm CpG probes. We introduce our proposed method in the following section and provide the analysis results of the DNHS data in Section 5.

## 3 Heterogeneous Mediation Analysis Method

In this section, we propose a heterogeneous mediator selection approach inspired by Tang et al. (2020) to achieve sub-population identification, mediator selection in each sub-population, and mediation effect estimation for heterogeneous data simultaneously. Statistically, to the best of our knowledge, this is the first work which considers heterogeneous mediation effects for high-dimensional potential mediators without pre-specifying subgroups. In addition, we do not assume the subgroup membership depending on observed covariates.

Moreover, to select mediators instead of variables in each sub-population, we propose a new mediation penalty which jointly penalizes the effect from the independent variable to a mediator (independent-mediator effect) and the effect from the mediator to the outcome (mediator-outcome effect). Essentially, the proposed mediation penalty encourages selection of mediators with large mediation effects. Before introducing the details of the proposed method, we first discuss related existing methods in the following subsection.

### 3.1 Existing Mediation Analysis methods

Traditional mediation analysis has been conducted via linear regression models (Baron and Kenny, 1986). As an extension, causal mediation analysis imposes “no unmeasured confounding” assumptions and defines direct and indirect effects under a counterfactual framework with potential outcomes (Rubin, 1974; Imai et al., 2010a; Robins and Greenland, 1992; Pearl, 2001).

Methods of multiple mediators have been developed in recent years (Serang et al., 2017; Boca et al., 2013; Imai et al., 2010b; Jirolon et al., 2020). To account for high-dimensional mediators, Zhao and Luo (2016) consider mediation pathway selection for a large number of causally dependent mediators, and present a sparse mediation model using a regularized structural equation model (SEM). In addition, Van Kesteren and Oberski (2018) develop an exploratory coordinate-wise mediation filter approach, and Zhang et al. (2016) propose a high-dimensional mediation analysis (HIMA) approach for DNA methylation. Moreover, Zhou et al. (2020) develop estimation and inference procedures for mediation effects under a high-dimensional linear mediation model. However, these methods are all under the homogeneous model framework.

To investigate heterogeneous mediation effects, Qin and Hong (2017) develop a weighting method to identify and estimate site-specific mediation effects, utilizing an inverse-probability-of-treatment weight (Rosenbaum, 1987) and ratio-of-mediator-probability weighting (Hong et al., 2015). This method assumes the heterogeneity caused by the variation of sites. However, the method is not applicable in general, since the potential mechanism resulting in sub-populations is usually unknown. Dyachenko and Allenby (2018) propose a Bayesian mixture model which combines likelihood functions based on two different outcome models to incorporate heterogeneity. Nevertheless, only a single mediator variable is considered and the mixture model requires a pre-specified number of subgroups.

### 3.2 Notations and Assumptions

In this subsection, we introduce notations and assumptions for the proposed method. Let *X*_*i*_ be an independent variable (e.g., treatment or exposure), ***Z***_*i*_ be a *r* × 1 vector of pre-treatment confounders (e.g., race or gender), ***M***_*i*_ = (*M*_*i*1_, …, *M*_*ip*_)^*T*^ be potential mediators, and *Y*_*i*_ be the outcome for the *i*-th subject (1 ≤ *i* ≤ *N*). Throughout the paper, we make the stable unit treatment value assumption (SUTVA) (Rubin, 1980); that is, the potential outcomes of one subject are unaffected by the assignment of treatments to other subjects, which is a standard assumption for causal inference. Without loss of generality, we also assume that the outcome and all the covariates are centered. We suppose that the entire population can be partitioned into *H* non-empty subgroups, where mediators and mediation effects within each subgroup are homogeneous. Denote the index set for subjects in the *h*-th subgroup by 𝒮(*h*) for *h* = 1, …, *H*.

### 3.3 Proposed Subgroup Linear Structural Equations Modeling Framework

We consider the heterogeneous mediation problem under the following subgroup linear structural equations modeling (LSEM) with *p* potential mediators and *r* pre-treatment confounders:

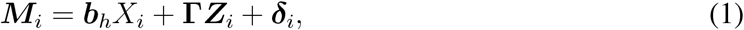

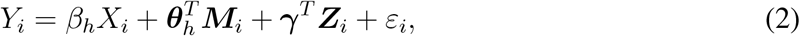

for subject *i* ∈ 𝒮(*h*), where ***b***_*h*_, ***δ***_*i*_, ***θ***_*h*_ ∈ ℝ^*p*^, **Γ** ∈ ℝ ^*p*×*r*^, and ***γ*** ∈ ℝ^*r*^. Here, *β*_*h*_ represents the direct effect from the independent variable *X*_*i*_ (e.g., trauma) to the outcome variable *Y*_*i*_ (e.g., PTSD) in the *h*-th subgroup, 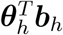 represents the joint mediation (indirect) effect of the treatment, ***b***_*h*_ is the parameter relating the treatment to the potential mediators, and ***δ***_*i*_ ∼ *N* (**0**, *Σ*) and *ε*_*i*_ ∼ *N* (0, *σ*^2^) are random errors, where ***δ***_*i*_ is independent of *X*_*i*_ and ***Z***_*i*_, and *ε*_*i*_ is independent of *X*_*i*_, ***Z***_*i*_, and ***M***_*i*_. We acknowledge that the LSEM imposes strong assumptions about the linearity and distribution of variables, indicating the results based on the proposed subgroup LSEM should be considered exploratory and instructive for generation of potential hypotheses.

We illustrate the relationship of variables in the LSEM for the *h*-th subgroup without the pretreatment confounders ***Z***_*i*_ in Figure 2. The arrow with *β*_*h*_ represents the direct effects from the independent variable *X*_*i*_ to the response *Y*_*i*_, while the arrow with ***θ***_*h*_ represents effects from mediators ***M***_*i*_ to the response *Y*_*i*_ in model (2). The direct effect *β*_*h*_ and indirect effects 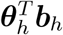 could vary across different subgroups.

**Figure 2:**
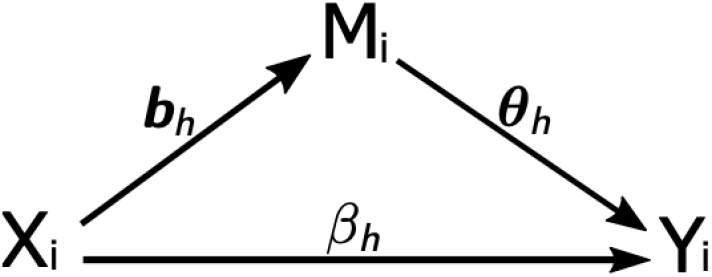
Mediation structure.

In this paper, we assume that the potential mediators are uncausally correlated (Jirolon et al., 2020). That is, the mediators could be conditionally dependent given the independent variable and observed pre-treatment confounders, but are not in any pre-specified causal order. For instance, it is possible that there exist an unmeasured covariate *U* affecting multiple mediators like two mediators *M*_1_ and *M*_2_ in 3(a), where *X* denotes the independent variable and *Y* denotes the outcome. Since *M*_1_ and *M*_2_ are not causally ordered, they are defined as “uncausally correlated”. In contrast, *M*_1_ and *M*_2_ in Figure 3(b) are causally ordered, that is, a change in *M*_2_ causes a change in *M*_1_.

**Figure 3:**
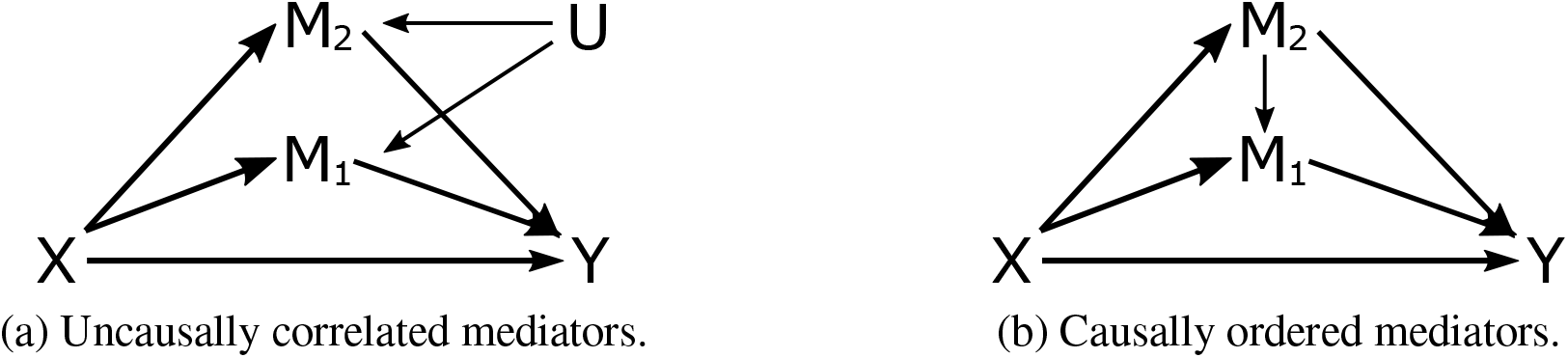
Different situations with two mediators.

In addition, we assume that *P* (*X*_*i*_ = *x* | ***Z***_*i*_ = ***z***) *>* 0 for all *x* and ***z*** if *X*_*i*_ is discrete, and that *f*_*X*_(*x* | ***Z***_*i*_ = ***z***) *>* 0 for all *x* and ***z*** if *X*_*i*_ is continuous, where *f*_*X*_ is the conditional density function of *X*_*i*_. We also suppose that the effects of pre-treatment confounders ***Z***_*i*_ on each mediator and the outcome are homogeneous across all the subgroups in (1) and (2). Moreover, we assume that the mediator models and the outcome model in (1) and (2) are sparse; that is, most true values in ***b***_*h*_ and ***θ***_*h*_ are zero for each *h*.

Under the proposed model and the assumptions, the sequential ignorability assumption in Jirolon et al. (2020) holds for each subgroup in the parametric model in (1) and (2). Moreover, the natural indirect effects ***δ***^*j*^(*t*) on the eighth page of Jirolon et al. (2020) for the *j*-th mediator is identical to *b*_*h,j*_*θ*_*h,j*_ for subjects from the *h*-th sub-population, where *b*_*h,j*_ and *θ*_*h,j*_ are the *j*-th element in ***b***_*h*_ and ***θ***_*h*_ in equations (1) and (2), respectively. In the following proposition, we have shown that parameters in (1) and (2) are identifiable, which implies that the indirect effects are identifiable under the parametric model. Thus we aim to estimate the causal quantity *b*_*h,j*_*θ*_*h,j*_ under the proposed model.

To state the proposition, we introduce some notations. Let ***C*** = (*C*_1_, …, *C*_*n*_)^*T*^ be a vector consisting of all subgroup labels, and **Θ** be a vector collecting all elements in **Γ, *γ*** and ***b***_*h*_, *β*_*h*_, ***θ***_*h*_ for *h* = 1, … *H*. Since the group-specific parameters, ***b***_*h*_, *β*_*h*_ and ***θ***_*h*_, are different across sub-groups, we consider the identifiability problem on the set 𝒜 = {(**Θ, *C***) : ***b***_*h*_ ≠ ***b***_*h*′_, *β*_*h*_ ≠ *β*_*h*′_, and ***b***_*h*_ ≠ ***b***_*h*′_ for any *h* ≠ *h*^′^ }.

#### Proposition 1.

*Under the non-degeneracy condition* **??**, *for any* (**Θ, *C***), (**Θ**^***′***^, ***C***^***′***^) ∈ 𝒜, *under the proposed model in equations (1) and (2), if the distribution of the observable variables satisfies f* (*X*_*i*_, ***M***_*i*_, *Y*_*i*_, ***Z***_*i*_; **Θ, *C***) = *f* (*X*_*i*_, ***M***_*i*_, *Y*_*i*_, ***Z***_*i*_; **Θ**^′^, ***C***^′^) *for* 1 ≤ *i* ≤ *N, then* (**Θ, *C***) *and* (**Θ**^′^, ***C***^′^) *are the same up to a permutation of subgroups*.

This proposition states that our model is parametrically identifiable up to a permutation of sub-groups. The proof of the Proposition is provided in Section S.1 of Supplementary Materials. Due to page limit, the non-degeneracy condition 1 is also in Section S.1 of Supplementary Materials.

To determine the subgroup membership of each subject, we evaluate how well each subject fits each sub-population by the loss function

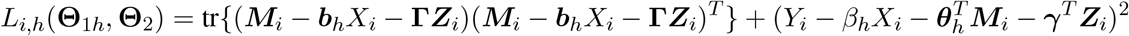

for the *i*-th subject and the *h*-th subpopulation, where 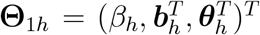 and **Θ**_2_ = (***γ*, Γ**^*T*^).

We propose to group subjects and identify the subgroup label of each subject through finding a smallest *L*_*i,h*_ for the *i*-th subject among all subgroups (*h* = 1, …, *H*), since a smaller loss function indicates better fitness and greater likelihood of the sample. Let **Θ**_1_ = (**Θ**_11_, …, **Θ**_1*H*_). Then, the loss function for all the subjects is

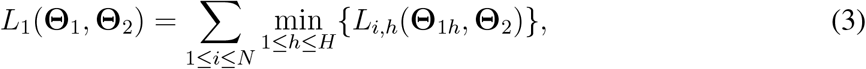

which is a sum of all within-cluster loss.

In the proposed method, we do not pre-specify or assume which variables determine the sub-group membership. Instead, these classes are identified in a completely data-driven manner, which is one of the key advantages of our method. We do not impose assumptions on which variables cause or determine the heterogeneity, since the heterogeneity of mediation effects could have complicated reasons. Essentially, analogous to a clustering algorithm, our model aggregates individuals into several classes where the individuals share similar mediation effects within each class but have different effects across classes. In addition, our proposed subgroup identification is different from that in the individualized-multi-directional method (IMDM) (Tang et al., 2021). We provide a comparison of the two methods in Section S.5.1 of the Supplementary Materials.

### 3.4 Mediation Regularization for Sparsity

In this subsection, we propose a new mediation penalty. As mentioned in Section 3.3, under the proposed subgroup LSEM in (1) and (2), the mediation effect of the *j*-th mediator in the *h*-th subgroup is *b*_*h,j*_*θ*_*h,j*_. To identify mediators with large mediation effects in each subgroup, we consider *b*_*h,j*_ and *θ*_*h,j*_ jointly for each 1 ≤ *j* ≤ *p*, and propose a two-dimensional joint mediation penalty

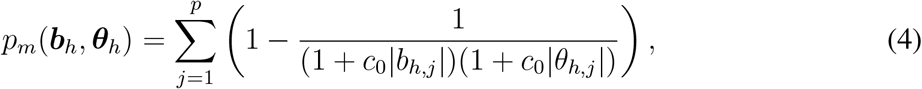

where *c*_0_ is a constant to adjust the shrinkage. We plot the mediation penalty with *p* = 1 and *c*_0_ = 0.5 in Figure 4. As shown in Figure 4, the penalty tends to shrink small values toward zero, and the shrinkage gradually levels off as |*θ*_*h*,1_| or |*b*_*h*,1_| increases.

**Figure 4:**
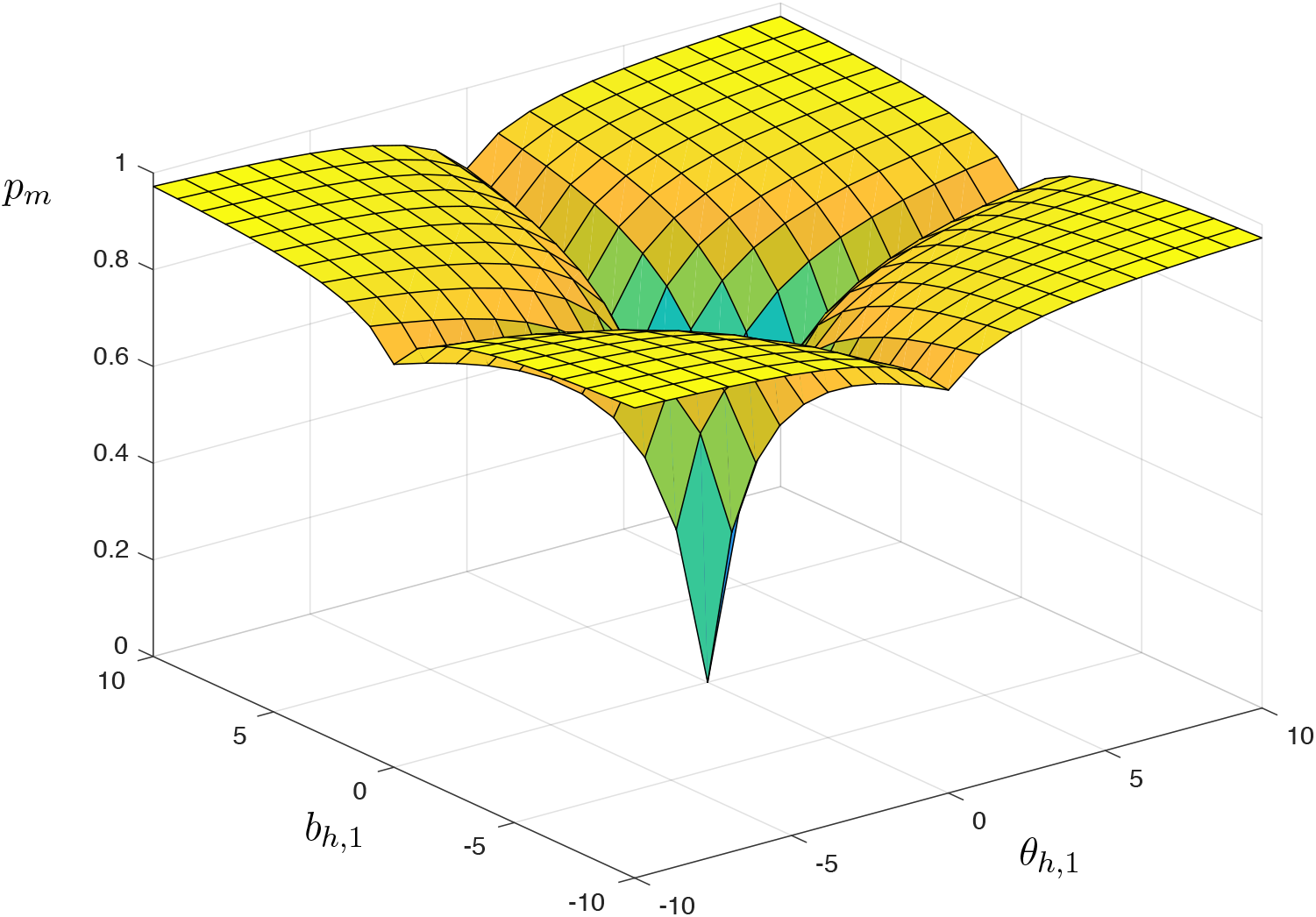
Mediation penalty with *p* = 1 and *c*_0_ = 0.5.

The high-dimensional mediation analysis proposed by Zhang et al. (2016) adopts the minimax concave penalty (MCP) (Zhang, 2010) for variable selection in the outcome model. Compared with the traditional Lasso (Tibshirani, 1996), MCP, and smoothly clipped absolute deviation (SCAD) penalties (Fan and Li, 2001), the proposed mediation penalty selects mediators instead of covariates in the sense that the *p*_*m*_(***b***_*h*_, ***θ***_*h*_) penalizes ***b***_*h*_ and ***θ***_*h*_ jointly rather than separately. Schaid and Sinnwell (2020) also jointly consider each pair of coefficients in the mediator and outcome model for each mediator using a group Lasso penalty. Compared to the group Lasso penalty, the proposed joint mediation penalty tends to be flat, and the corresponding shrinkage gradually levels off as the coefficients in each pair increase; which can relax the rate of penalization for large coefficients and large mediation effects. Thus, the proposed penalty could reduce the bias due to joint shrinkages.

Intuitively, we need a relatively small shrinkage for *b*_*h,j*_ when *θ*_*h,j*_ is large; otherwise it is hard to select the *j*-th mediator when *b*_*h,j*_ is small but *θ*_*h,j*_ and the mediation effect *b*_*h,j*_*θ*_*h,j*_ are large. However, this property does not hold for penalty functions such as |*θ*_*h,j*_| + |*b*_*h,j*_| and the pathway Lasso penalty in Zhao and Luo (2016) penalizing ***b***_*h*_ and ***θ***_*h*_ jointly via |*θ*_*h,j*_*b*_*h,j*_| (1 ≤ *j* ≤ *p*). Specifically, with a penalty of |*θ*_*h,j*_| + |*b*_*h,j*_|, the shrinkage of *b*_*h,j*_ does not depend on *θ*_*h,j*_; and with a penalty of |*θ*_*h,j*_*b*_*h,j*_|, the shrinkage on *b*_*h,j*_ increases as *θ*_*h,j*_ increases. In contrast, our joint mediation penalty in (4) has the desired property since the shrinkage on *b*_*h,j*_ in *p*_*m*_(***b***_*h*_, ***θ***_*h*_) gradually levels off as *θ*_*h,j*_ increases and vice versa, as shown in Figure 4. On the other hand, the proposed mediation penalty will not induce a strong shrinkage for *θ*_*h,j*_ if *b*_*h,j*_ = 0. In Figure 4, even if *b*_*h*,1_ = 0, the rate of penalization on *θ*_*h*,1_ will still decrease as *θ*_*h*,1_ itself increases, which is analog to the SCAD penalty imposed on a single coefficient *θ*_*h*,1_.

### 3.5 Fusion Penalty for Cross-group Information

In general, it is possible that not every mediator has different mediation effects across subgroups. We use the following between-group fused Lasso penalty (Tibshirani et al., 2005)

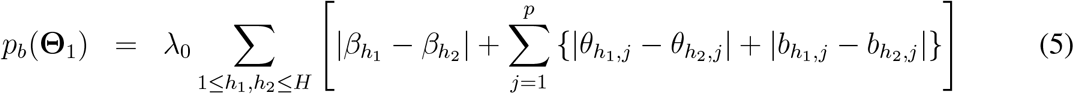

to shrink similar between-group effects together, where *λ*_0_ is a tuning parameter. Specifically, the between-group penalty *p*_*b*_(**Θ**_1_) encourages mediators to share the same parameter across different subgroups when the corresponding effects are similar. In this way, we can borrow information across subgroups in estimating the mediation effects.

Consequently, the objective function of the proposed method is

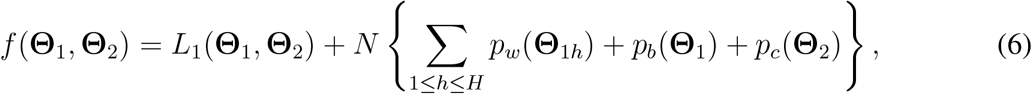

where

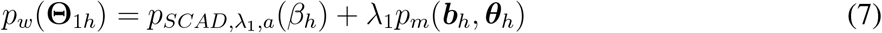

is a within-group penalty, 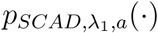 is the SCAD penalty with tuning parameters *λ*_1_ and 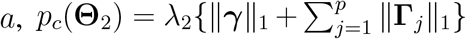 is a *l*_1_ penalty with tuning parameter *λ*_2_ for pre-treatment confounders to avoid over-fitting, and **Γ**_*j*_ denotes the *j*-th row of **Γ**. Through minimizing *f* (**Θ**_1_, **Θ**_2_) in (6), we not only incorporate the heterogeneity of mediation effects in subjects, but also combine cross-group information for similar effects. Moreover, in the objective function *f* (**Θ**_1_, **Θ**_2_), we penalize not only ***θ***_*h*_ but also the direct effect *β*_*h*_ in the outcome model in equation (2) since otherwise the estimation obtained by minimizing the objective function could falsely transfer the effects of mediators ***M***_*i*_ to the effect of *X*_*i*_. In the following subsection, we provide an algorithm to obtain the minimizer of the objective function.

### 3.6 Implementation

In this subsection, we propose an effective algorithm to minimize the objective function in equation (6). Note that our objective function is non-convex, since the within-group penalty *p*_*w*_ in (7) is not convex and the loss function in (3) involves minimization. To tackle this challenge, we decompose our objective function as a difference of two convex functions and solve the optimization problem based on the difference of convex (DC) algorithm (Le Thi Hoai and Tao, 1997; Shen et al., 2012), which is shown to converge to a stationary point under regularity conditions (Abbaszadehpeivasti et al., 2021). Also, we use a smooth approximation of the between-group fused Lasso penalty.

Specifically, we rewrite the objective function as a difference of two functions

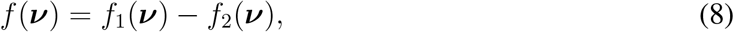

where ***ν*** is a vector consisting of all parameters ***b***_*h*_, *β*_*h*_, ***θ***_*h*_, ***γ*, Γ** (for *h* = 1, …, *H*) in models (1) and (2),

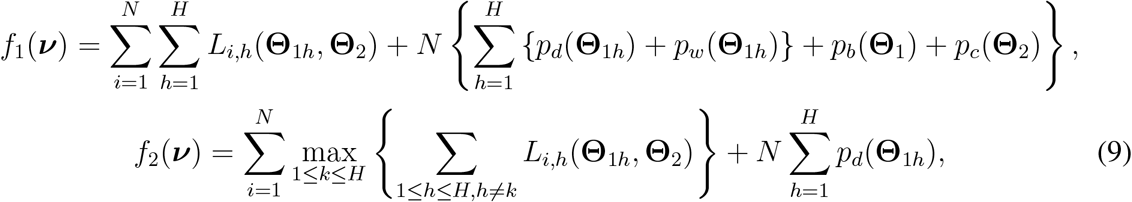

and 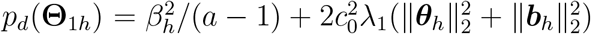. It can be shown that *p*_*d*_(**Θ**_1*h*_) + *p*_*w*_(**Θ**_1*h*_) is convex for each 1 ≤ *h* ≤ *H*, since the corresponding Hessian matrix is block-diagonal and each diagonal block is positive-definite. This implies that *f*_1_(***ν***) is a convex function. In addition, since the maximum of convex functions is still convex, *f*_2_(***ν***) is also a convex function.

Based on this DC decomposition on *f*, we iteratively construct a sequence of convex approximations for *f* (***ν***). We first calculate the subdifferential of *f*_2_(***ν***) in the following:

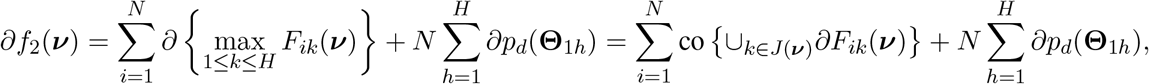

where *F*_*ik*_(***ν***) = 1≤*h*≤*H,h*≠*k L*_*i,h*_(**Θ**_1*h*_, **Θ**_2_), *J* (***ν***) = {1 ≤ *k* ≤ *H* : *F*_*ik*_(***ν***) = max_1≤*k*≤*H*_ *F*_*ik*_(***ν***)}, and “co” stands for the convex hull. At the *m*-th iteration, given the previous estimate ***ν***^(*m*−1)^, we replace *f*_2_(***ν***) in (8) by its affine minorization: 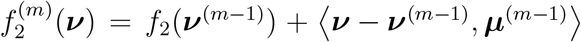, where 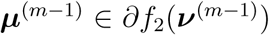. Then, 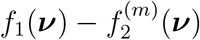 is an upper convex approximating function for *f* (***ν***) at the *m*-th iteration. Through this, we convert the non-convex objective function into a convex relaxation via a tangent approximation of *f*_2_(***ν***).

In addition, since the between-group fused Lasso *p*_*b*_(**Θ**_1_) in *f*_1_(***ν***) is non-smooth and non-separable, we approximate it by a smooth function. Specifically, we reformulate the fused Lasso as 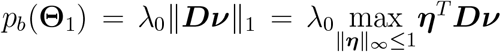, where ***D*** is a difference operator corresponding to the differences in *p*_*b*_(**Θ**_1_). That is, the *p*_*b*_(**Θ**_1_) is equivalent to the maximum of the maximization problem for ***η***^*T*^ ***Dν***. Hence, we let 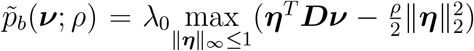, where *ρ* is a positive smoothing parameter. This function 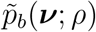 approximates *p*_*b*_(***ν***) as *ρ* → 0 (Nesterov, 2005). Let ***η***^*^ = *S*(***Dν****/ρ*), where

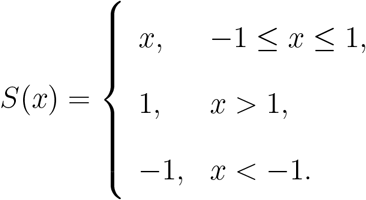

Then we have 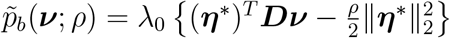, which is convex and differentiable in ***ν*** (Chen et al., 2012). In our implementation, we choose *ρ* = 10^−4^ following Chen et al. (2012).

Consequently, at the *m*-th iteration, we replace the *f*_2_(***ν***) and *p*_*b*_(***ν***) in (8) by 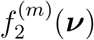 and 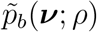, respectively, and obtain

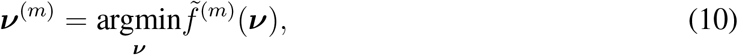

where

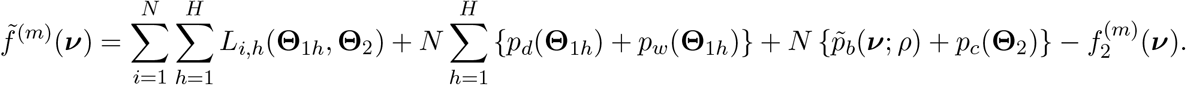

We solve the minimization problem in equation (10) through the gradient descent algorithm (Curry, 1944) with the backtracking line search (Shi, 2004; Stanimirović and Miladinović, 2010) for the step size. The above algorithm can be summarized in Algorithm 1. We also provide an expanded version of this algorithm in Section S.2 of the Supplementary Materials.

#### Algorithm 1

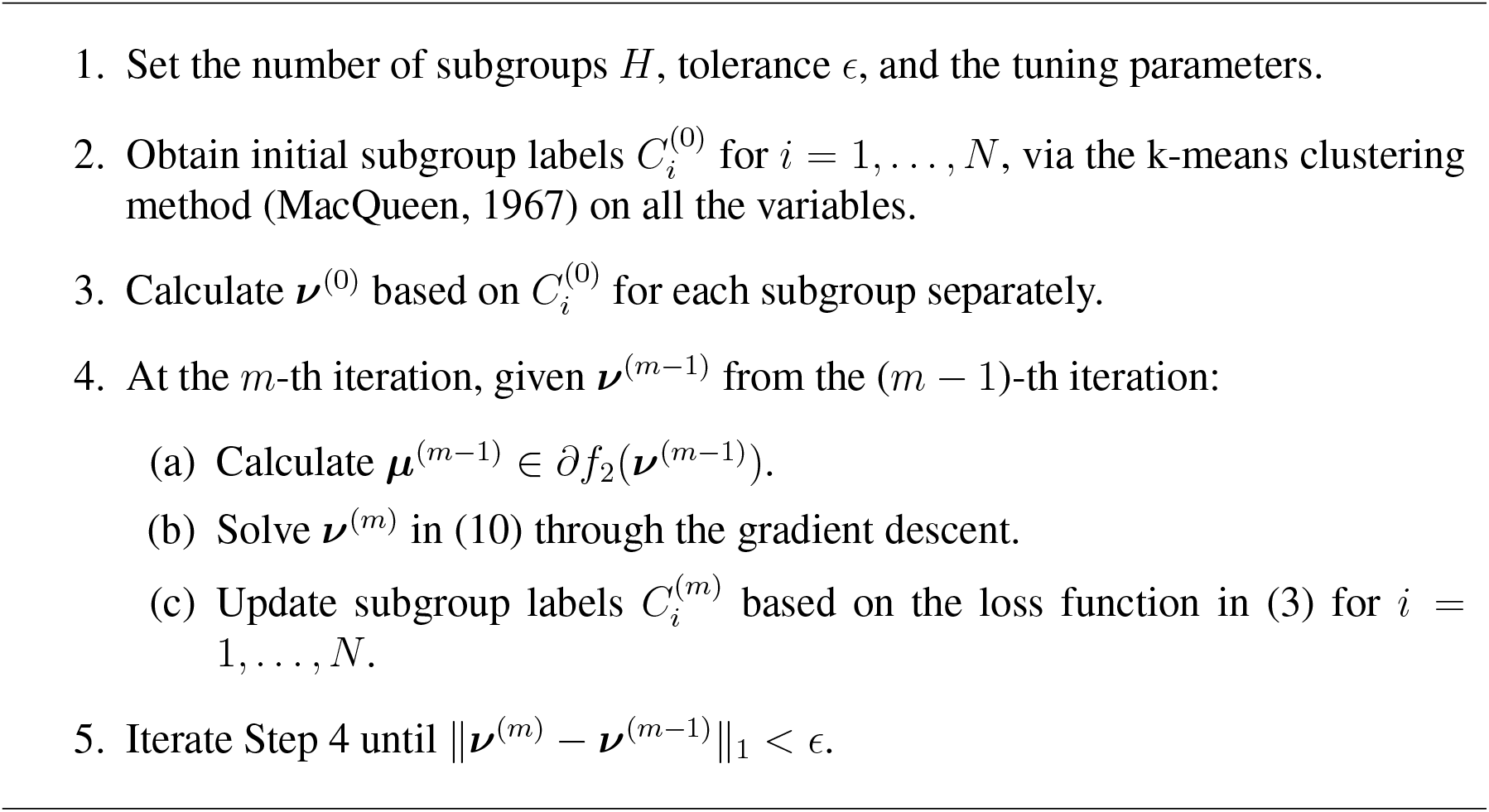

To determine the tuning parameters and the number of subgroups *H*, we propose the following Bayesian information criterion (BIC) type criterion:

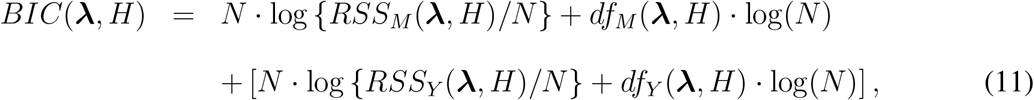

where *df*_*M*_ (***λ***, *H*) and *df*_*Y*_ (***λ***, *H*) are numbers of non-zero estimated coefficients in mediator models and the outcome model, respectively, 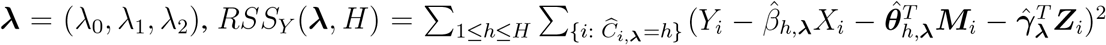, and 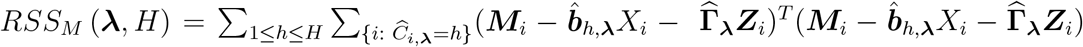. Here, 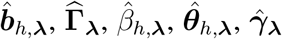, and 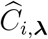 are the estimates of ***b***_*h*_, **Γ**, *β*_*h*_, ***θ***_*h*_, ***γ***, and *C*_*i*_, respectively, based on *H* subgroups and tuning parameters ***λ***. Recall that the between-group fused Lasso penalty *p*_*b*_(**Θ**_1_) in (5) encourages shared parameters for similar effects across different subgroups. In *df*_*M*_ and *df*_*Y*_, the shared parameters are counted without multiplicity.

We select the optimal tuning parameters and the optimal number of subgroups through minimizing *BIC*(***λ***, *H*), incorporating information from both the mediator models and the outcome model. In the implementation for the following sections, we mainly tune *λ*_0_, *λ*_1_, and *H* using a grid search to minimize the BIC. We do not tune *λ*_2_ since it is for penalization of the pre-treatment confounders, which are not involved in our simulations and real data application.

## 4 Simulated Data Experiments

In this section, we investigate the performance of the proposed method compared with existing homogeneous mediation methods via simulation studies. We simulate data following models in (1) and (2) with *r* = 0. The proposed method is implemented based on Algorithm 1 with *c*_0_ = 10 and *a* = 3.7. We apply the “HIMA” package (https://cran.r-project.org/web/packages/HIMA/index.html) in R to implement the high-dimensional mediation analysis (HIMA) method (Zhang et al., 2016) for comparison, which is a homogeneous mediation approach. Our results are summarized based on 100 replications.

To evaluate the performance of each method, we calculate the average of all individuals’ mediator false negative rates (FN) and the average of all individuals’ mediator false positive rates (FP) for mediator selection as follows:

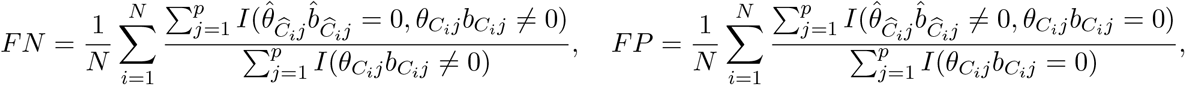

where 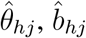, and 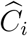 are estimators of *θ*_*hj*_, *b*_*hj*_, and *C*_*i*_, respectively. Specifically, the FN and FP represent proportions of unselected true mediators and selected noises, respectively. A method with smaller FN+FP selects more accurate mediators. In practice, the FP and FN may have different costs. For example, in scenarios such as a pregnancy test or a COVID-19 test, FN costs much more than FP. However, in other scenarios such as criminal conviction or identifying spam emails, FP costs much more than FN. Given specific application context and background information, we may assign different weights to FP and FN, respectively. Since simulation studies do not involve any real situations, we treat them equally and just use FN+FP as an evaluation criterion.

We also evaluate each method via the mean-squared-errors (MSE) of mediation effects in an average of all individuals as 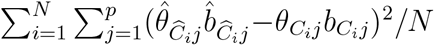. For the proposed method, we also report the proportion of replications where the number of subgroups is correctly selected via the *BIC*(***λ***, *H*) in (11), which we refer to as the correct rate of subgroup number selection. Moreover, we compute the misclassification rate of subjects based on the proposed method for evaluation of subgroup identification.

We consider the following three settings, where the non-zero coefficients in ***b***_*h*_ and ***θ***_*h*_ share the same signal strengths *b*_*hs*_ and *θ*_*hs*_, respectively, for 1 ≤ *h* ≤ *H*. For the sensitivity of the proposed approach to misspecification, we investigate situations without heterogeneity in the first setting, which involves a homogeneous underlying true model with only one sub-population. In contrast, Settings 2 and 3 assume heterogeneous true models with two sub-populations. In addition, we consider high-dimensional situations in Setting 3.

### Setting 1.

*Let N* = 200, *H* = 1, *n*_1_ = 200, *and p* = 30. *True coefficients in the model are illustrated in Figure 5 with β*_1_ = 0.5, *b*_1*s*_ = 1, *and θ*_1*s*_ = 0.2, 0.3, *or* 0.4. *As shown in Figure 5, we have four true mediators with mediation effects θ*_1*s*_. *In addition, we generate X*_*i*_ *and ε*_*i*_ *from a standard normal distribution and* ***δ***_*i*_ ∼ *N* (**0, *I***_*p*×*p*_) *for each i* = 1, …, *N*.

**Figure 5:**
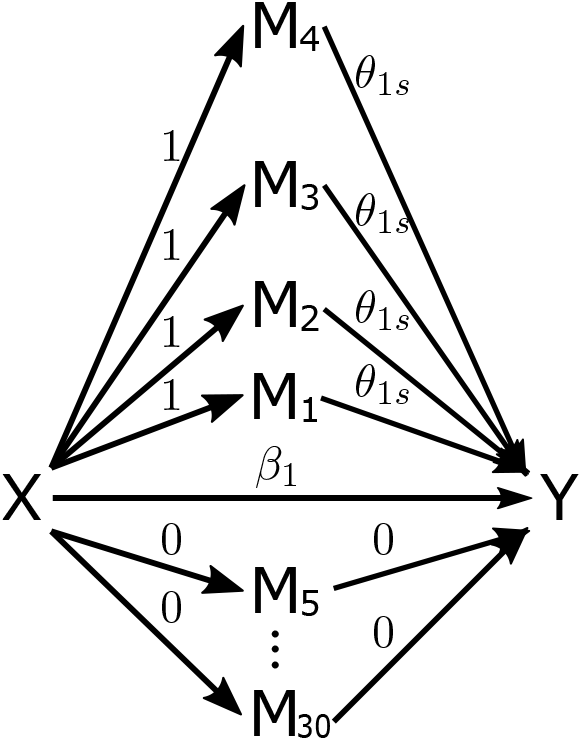
True coefficients for the homogeneous Setting 1. The “X” denotes the independent variable, “M_*i*_” denotes the *i*-th mediator, and “Y” denotes the dependent variable. The value above each arrow represents the true coefficient for the corresponding effect.

### Setting 2.

*We proceed similarly as in Setting 1 except that H* = 2, *n*_1_ = 50, *and n*_2_ = 150. *True coefficients in the model are illustrated in Figure 6 with β*_1_ = 0.5, *b*_1*s*_ = 1, *β*_2_ = −0.5, *b*_2*s*_ = −1, *θ*_1*s*_ = 0.5, 1, *or* 4 *and θ*_2*s*_ = −0.5, −1, *or* −4. *As shown in Figure 6, we have three true mediators in each subgroup*.

**Figure 6:**
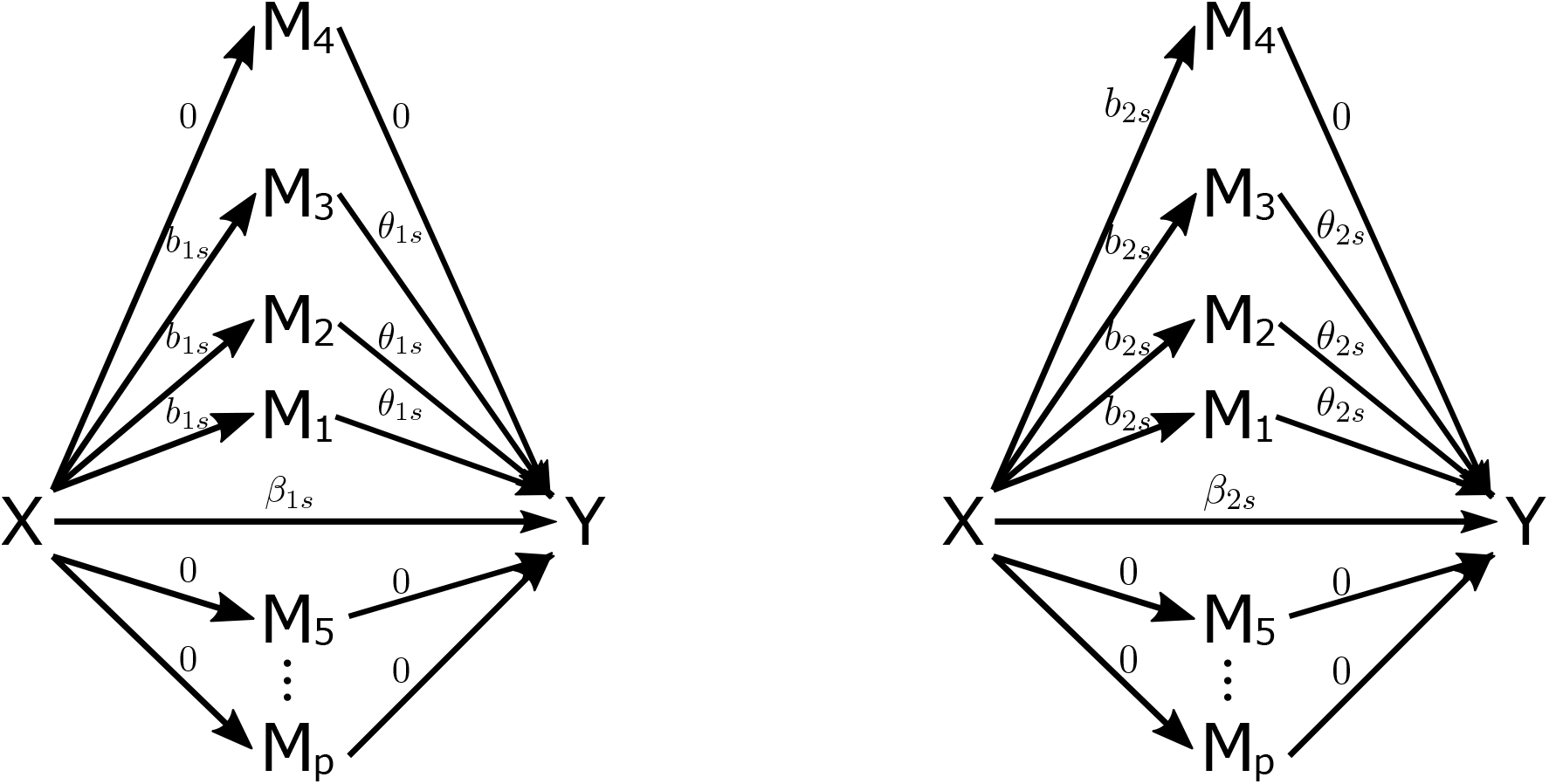
True coefficients for the heterogeneous Settings 2 and 3 with two sub-populations (left and right). The *p* is 30 and 150 under Settings 2 and 3, respectively. The “X” denotes the independent variable, “M_*i*_” denotes the *i*-th mediator, and “Y” denotes the dependent variable. The value above each arrow represents the true coefficient for the corresponding effect.

### Setting 3.

*We investigate a high-dimensional case proceeding similarly as in Setting 2 except that N* = 100, *n*_1_ = 30, *n*_2_ = 70, *p* = 150, *β*_1_ = 1, *β*_2_ = −1, *θ*_1*s*_ = 0.5, 0.8, 1, *or* 4 *and θ*_2*s*_ = −0.5, −0.8, −1, *or* − 4. *The covariance matrix of* ***δ***_*i*_ *has an autoregressive structure of order* 1, *that is, AR(1), with diagonal* 1 *and off-diagonal parameter ρ*.

Tables 2–4 provide the results of the proposed method and the HIMA method, and show that the proposed method produces smaller overall FN+FP totals than the HIMA method across all the settings, indicating that the proposed method selects mediators more accurately. Moreover, the proposed method produces smaller MSE of mediation effects, implying that the proposed method is also more effective in estimation of mediation effects.

**Table 2:** FN, FP, FN+FP, and MSE under Setting 1. “HIMA” stands for the high-dimensional mediation analysis method. The “*θ*_1*s*_” represents the signal strength in ***θ***_1_.

| $\theta_{1s}$ | Method | FN | FP | FN+FP | MSE |
| --- | --- | --- | --- | --- | --- |
| 0.2 | **Proposed** | 0.046 | 0.002 | 0.048 | 0.002 |
|  | HIMA | 0.570 | 0.000 | 0.570 | 0.003 |
| 0.3 | **Proposed** | 0.000 | 0.000 | 0.000 | 0.002 |
|  | HIMA | 0.100 | 0.000 | 0.100 | 0.002 |
| 0.4 | **Proposed** | 0.000 | 0.000 | 0.000 | 0.002 |
|  | HIMA | 0.005 | 0.000 | 0.005 | 0.001 |

In particular, under the homogeneous Setting 1, the proposed method still outperforms the homogeneous HIMA method. For example, when *θ*_1*s*_ = 0.2, the FN+FP of the proposed is only 8.8% of that of the HIMA as shown in Table 2. This is likely due to the advantage of the proposed mediation penalty, and that the proposed method correctly identifies the number of subgroups in most situations as shown in Table 5. Moreover, the proposed method falsely un-selects 4.6% of true mediators and falsely selects just 0.2% of noises (variables that are not mediators), even when the mediation effect of each true mediator is as small as 0.2. In addition, under this situation, the MSE of mediation effects is only 0.002, indicating that the proposed method performs consistently well even when there is no heterogeneity and the effect size is small.

In addition, the proposed method also performs much better than the HIMA when there are two subgroups with opposite mediation effects in Settings 2 and 3, that is, with positive mediation effects in one subgroup and negative mediation effects in the other. In this case, the homogeneous HIMA method usually fails to identify true mediators. For instance, the FN of the HIMA is as high as 0.957 when (*θ*_1*s*_, *θ*_2*s*_) = (0.5, −0.5) as shown in Table 3, while the corresponding FN of the proposed method is only 0.031.

**Table 3:** FN, FP, FN+FP, and MSE under Setting 2. “HIMA” stands for the high-dimensional mediation analysis method. The “*θ*_1*s*_” and “*θ*_2*s*_” represent the signal strength in ***θ***_1_ and ***θ***_2_, respectively.

| $(\theta_{1s}, \theta_{2s})$ | Method | FN | FP | FN+FP | MSE |
| --- | --- | --- | --- | --- | --- |
| (0.5, -0.5) | **Proposed** | 0.031 | 0.007 | 0.038 | 0.004 |
|  | HIMA | 0.963 | 0.004 | 0.968 | 0.025 |
| (1, -1) | **Proposed** | 0.002 | 0.000 | 0.002 | 0.006 |
|  | HIMA | 0.943 | 0.003 | 0.946 | 0.099 |
| (4, -4) | **Proposed** | 0.001 | 0.000 | 0.001 | 0.058 |
|  | HIMA | 0.790 | 0.002 | 0.792 | 1.514 |

Moreover, we explore high-dimensional scenarios in Setting 3 to mimic the case of the DNHS data. In this setting, we also consider the situations where the error terms in the mediator models in (1) are correlated, that is, the correlations among mediators may not just come from the independent variable. In all the high-dimensional cases, the proposed method produces smaller FN+FP and smaller MSE than the HIMA method illustrated in Table 4. For example, when (*θ*_1*s*_, *θ*_2*s*_) = (4, −4) and *ρ* = 0.2, the FN+FP of the proposed method is only 16.7% of that of the HIMA method, and the MSE is only 29.8% of that of the HIMA.

**Table 4:**
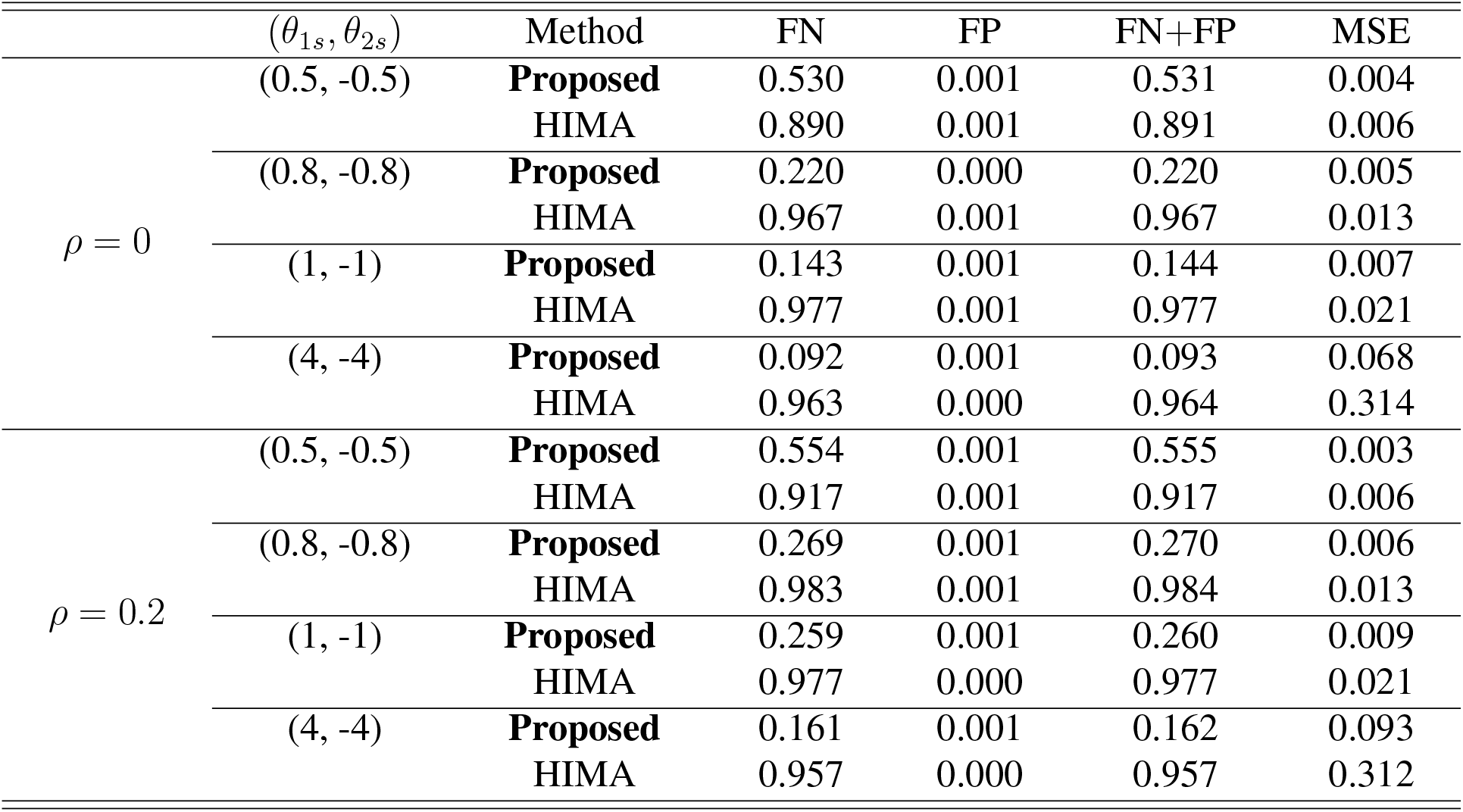
FN, FP, FN+FP, and MSE under Setting 3. “HIMA” stands for the high-dimensional mediation analysis method. The “*θ*_1*s*_” and “*θ*_2*s*_” represent the signal strength in ***θ***_1_ and ***θ***_2_, respectively, and “*ρ*” is a correlation parameter.

In Table 5, we provide the correct rates of subgroup number selection and the misclassification rates under various settings. We observe that the proposed method correctly determines the number of subgroups in most situations, especially in Settings 1 and 2 where there are more samples. In addition, the proposed method groups most subjects correctly across different settings due to the low misclassification rates.

**Table 5:** Correct rate of subgroup number selection and misclassification rate for different settings. “Correct rate of subgroup number selection” represents the proportion of replications where the number of subgroups is correctly selected via the proposed method. “Misclassification rate” is the average proportion of subjects who are mis-classified by the proposed method.

| | $(\theta_{1s}, \theta_{2s})$ | Correct rate of subgroup number selection | Misclassification rate |
| --- | --- | --- | --- |
| Setting 1 | (0.2, -) | 0.76 | – |
|  | (0.3, -) | 1.00 | – |
|  | (0.4, -) | 1.00 | – |
| Setting 2 | (0.5, -0.5) | 0.99 | 0.13 |
|  | (1, -1) | 1.00 | 0.11 |
|  | (4, -4) | 0.92 | 0.10 |
| Setting 3<br>$\rho = 0$ | (0.5, -0.5) | 0.66 | 0.17 |
|  | (0.8, -0.8) | 0.68 | 0.14 |
|  | (1, -1) | 0.72 | 0.13 |
|  | (4, -4) | 0.62 | 0.12 |
| Setting 3<br>$\rho = 0.2$ | (0.5, -0.5) | 0.59 | 0.17 |
|  | (0.8, -0.8) | 0.69 | 0.15 |
|  | (1, -1) | 0.62 | 0.15 |
|  | (4, -4) | 0.66 | 0.14 |

In summary, our simulation studies show that the proposed method achieves higher mediator selection accuracy and mediation effect estimation accuracy than the existing homogeneous mediation method across all the settings. One reason is that the proposed method adopts the mediation penalty in (4) which considers effects in mediator models and outcome models jointly, and encourages selection of mediators with large mediation effects. In addition, the proposed method allows heterogeneity among subjects, and thus can identify mediators with heterogeneous mediation effects, which is especially powerful for mediators with opposite effects in different subgroups. We investigate more simulations for larger-scale cases, with different initial values, various coefficients, moderators, and under non-normality of the independent variable in Sections S.3.1, S.3.2, S.3.4–S.3.6 of Supplementary Materials, respectively.

## 5 DNHS Case Study

In this section, we investigate how DNA methylation mediates the effects of traumatic experiences on development of PTSD based on the DNHS data. Specifically, we apply the proposed method to study the mediation effects of DNAm variation of GRRN genes on PTSD symptom severity. In this study, we use the baseline wave of the DNHS data for our mediation analysis. We treat the total number of trauma exposures as an independent variable, DNAm CpG probes that are significantly correlated with the GRRN genes as potential mediators, and the average PCL-C score as the outcome variable. There are 125 subjects and 144 selected DNAm CpG probes after screening. Our main objective is to identify key DNAm CpG probes from all the potential mediators.

To evaluate the performance of the proposed method compared with existing methods, we randomly split the data into a training set (90% of all samples) and a testing set (10% of all samples) for 100 times. The training sets in the 100 replications are repeated random subsamples (90%) of the complete data. For the proposed method, to identify the subgroup label of the *i*-th subject in the testing set, we calculate the average prediction error of mediators 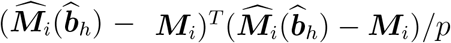, where 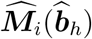 is the predicted value for ***M***_*i*_ based on estimated parameters 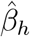 and 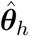 in the *h*-th subgroup. Then the *i*-th subject is labeled with a specific subgroup that minimizes 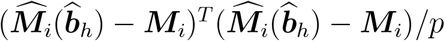. For each method, we train the model on the training data, and calculate the prediction root-mean-squared errors (PRMSEs) for the mediators and the outcome variable in the testing set; that is, 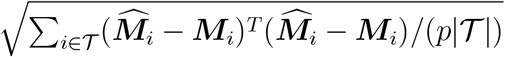 and 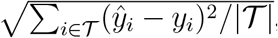, respectively, where 𝒯 denotes the index set of subjects in the testing set and |𝒯| is the testing sample size.

We provide mean PRMSE for each method based on 100 replications in Table 6. The proposed method produces much smaller prediction errors for both the mediators and the outcome variable compared to the existing high-dimensional mediation analysis (HIMA) method, indicating that the proposed method is more accurate in terms of prediction. Note that, in both methods, we use all the potential mediators to calculate the prediction errors of mediators, and then take an average of the errors over the mediators.

**Table 6:** The DNHS data results by the proposed method and the high-dimensional mediation analysis (HIMA) method. Here “NS” represents the mean number of selected mediators. The “Mediator” represents average prediction error for mediators, and “Outcome” represents average prediction error for the outcome variable based on 100 replications.

| Method | NS | Mediator | Outcome |
| --- | --- | --- | --- |
| **Proposed** | 6 | 0.455 | 1.235 |
| HIMA | 0 | 4.309 | 2.774 |

For the subgroup identification in the DNHS data, we apply the proposed method to all the samples. The proposed method selects four DNAm probes, and identifies three subgroups consisting of 60, 26, 39 subjects, respectively. We provide the estimated coefficients of the four DNAm probes in the three subgroups in Figure 7. Although the size of some estimated coefficients are small, the estimated mediation effect size here is comparable to the findings from other studies for DNA methylation (Tobi et al., 2018), and thus the mediation effects are still non-ignorable. For example, the mediation effect from trauma to PCL score through cg01277438 is 0.02 × 0.94 ≈ 0.02 for the subgroup in Figure 7(c). Note that the range of the trauma measurement is from 0 to 14 with mean 4.0. If the trauma variable takes its average value 4.0, the influence of this trauma value on PCL score through the mediator cg01277438 is about 0.08. Moreover, if the trauma variable takes its maximum value 14, the influence of this trauma value on the PCL score through the mediator cg01277438 is about 0.28, which is 25% of the standard deviation 1.1 of the PCL score.

**Figure 7:**
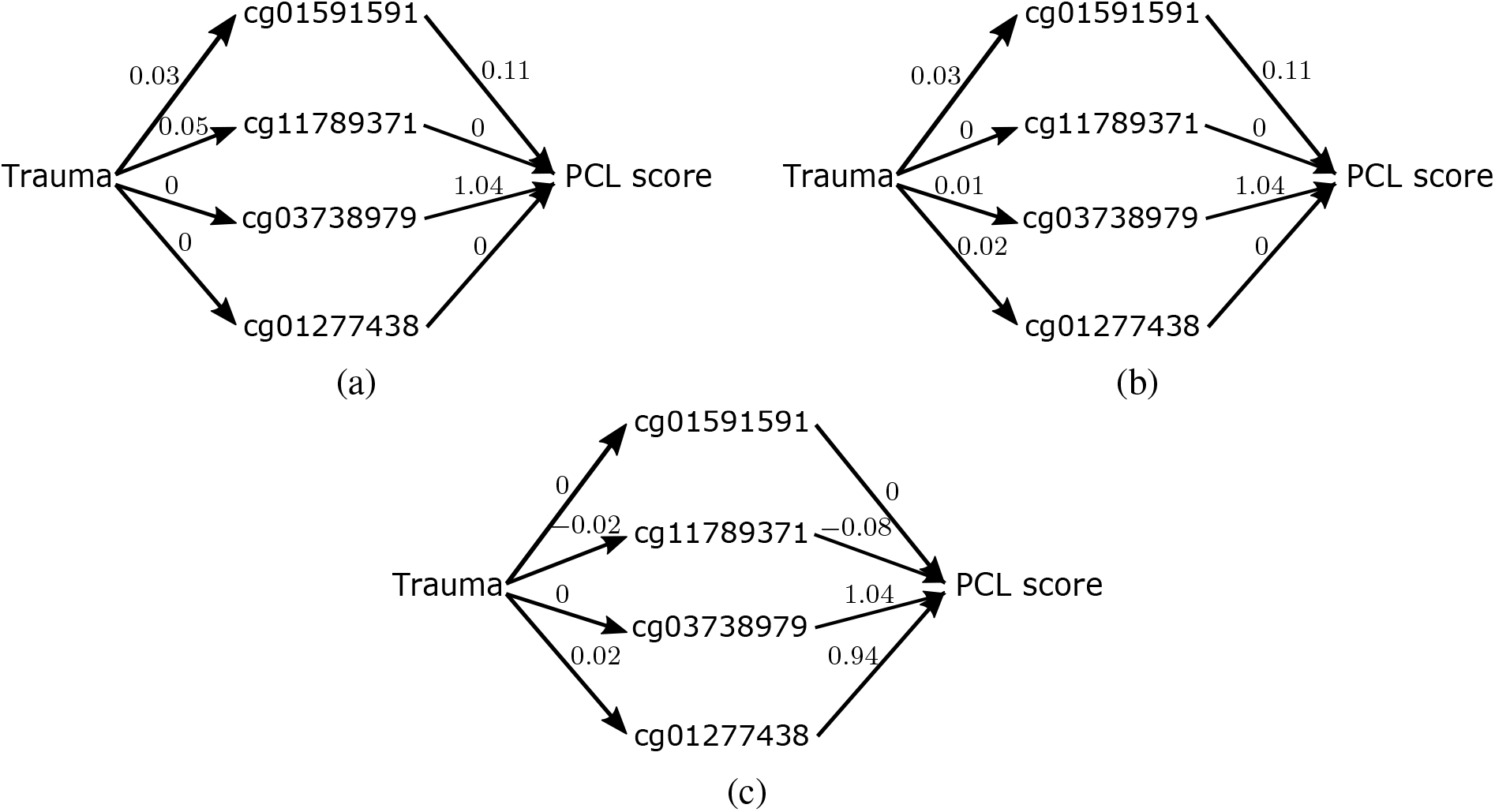
Estimated coefficients for the selected four mediators in the three subgroups identified by the proposed method based on all samples.

Specifically, the four DNAm probes identified by the proposed method correspond to *NFATC1, HSP90AA1, SMARCA4*, and *CREBBP* genes, among which *HSP90AA1, SMARCA4*, and *NFATC1* are indeed related to PTSD based on existing literature (Criado-Marrero et al., 2018; Breen et al., 2019; Kim et al., 2019a; Raabe and Spengler, 2013; Kim et al., 2019a; Kuan et al., 2017). In addition, we apply the HIMA method to each of the three subgroups on the four mediators. At a 0.05 significance level, the HIMA method shows that three (“cg11789371”, “cg03738979”, “cg01277438”) of the four selected mediators are significant, after controlling the false discovery rate (FDR) via the Benjamini–Hochberg procedure (Benjamini and Hochberg, 1995). However, the HIMA method cannot identify any mediator when all potential mediators and samples are used. This confirms that the subgroups and mediators selected by the proposed method are useful in scientific findings.

Furthermore, we find common patterns in subgroup identification across the analyses of the 100 random split datasets and the whole dataset, which are provided in Section S.4.1 of Supplementary Materials. Also, we investigate the racial make-up of the three identified subgroups. The results are provided in Table 13 in Section S.4.2 of Supplementary Materials, showing that the subjects are not grouped based on race since each subgroup contains both AAs and European Americans.

In summary, the proposed method produces smaller prediction errors than the homogeneous mediation method. Moreover, the proposed method identifies important mediators which cannot be detected by existing methods. In addition, our method shows heterogeneous mediation among subjects in the DNHS data.

## 6 Discussion

Our main contribution is to advance the understanding of the underlying mechanism of PTSD using the DNHS data. Specifically, we conduct a mediation analysis for DNA methylation on the relationship between traumatic events and PTSD symptoms based on the DNHS data. The identification of DNAm mediators presents new statistical challenges due to the heterogeneous nature of PTSD among patients and the high-dimensional structure of the DNAm data. To address these challenges, we develop a heterogeneous mediation model with multiple mediators, incorporating heterogeneity among subjects. Moreover, we propose a novel mediation penalty to incorporate effects in the mediator models and the outcome model jointly for high-dimensional data. Our numerical studies show that the proposed method selects true mediators more accurately than the existing homogeneous high-dimensional mediation analysis method.

In the DNHS case study, we identify meaningful DNAm mediators using the proposed method, which has important impact in practice since it would advance development of new personalized treatments for PTSD. In fact, recent studies have shown that successful PTSD treatments are reflected by significant DNAm changes (Vinkers et al., 2019; Yehuda et al., 2013). Our finding could also suggest future biomedical research on the selected mediators and corresponding genes for further biological verification. In particular, the selected DNAm CpG probes correspond to genes such as *HSP90AA1, SMARCA4*, and *NFATC1*, which have been reported in literature that they are associated with the PTSD (Criado-Marrero et al., 2018; Breen et al., 2019; Kim et al., 2019a; Raabe and Spengler, 2013; Kim et al., 2019a; Kuan et al., 2017).

In addition, the subgroup identification by the proposed method for the DNHS dataset with predominantly African-American subjects indicates potential heterogeneity in the underlying DNAm profiles which mediate risk for PTSD. This is an important discovery, as it could help us to uncover important genetic complexities which have been ignored for treatment and prevention of this debilitating mental disorder.

In this article, we mainly consider the detection of heterogeneous mediators and estimation of corresponding mediation effects. We have not developed statistical inference for the heterogeneous mediation effects, which is a limitation of this paper. It would be of great interest to investigate the statistical inference on these mediation effects in the future; for example, constructing de-biased estimators of sub-population mediation effects for confidence intervals or hypothesis testing. A de-biasing procedure can also reduce the bias for mediation effects incurred from regularization. One possible way is to control the bias of estimators for parameters in both mediator and outcome models, and then use multiplication of each pair of de-biased estimates to estimate the mediation effects of the corresponding mediator.

Moreover, the proposed model in equations (1) and (2) implies that there is no unmeasured confounders. However, to our best knowledge, it is unclear whether there are any common factors impacting both methylation and PTSD symptoms, as the exact biological processes underlying the development of PTSD remain elusive, which is a study limitation given the current state of science. We provide more discussion on the proposed model regarding exposure–mediator interactions, latent subpopulations, and moderated mediation and mediated moderation in Sections S.5.2–S.5.4 of Supplementary Materials, respectively.

Furthermore, our use of cross-sectional rather than longitudinal methylation measurements in the DNHS data application can be considered a study limitation, since the mediation assumption cannot be verified in cross-sectional data. However, it is consistent with the majority of existing epigenetic work relevant to PTSD published to date. In fact, many methylation studies are actually cross-sectional (Christiansen et al., 2021; Nakatochi et al., 2017; Alghanim et al., 2017; Joehanes et al., 2016; Young et al., 2016; King et al., 2014), and the majority of epigenetic PTSD studies to date have used a similar study design (Sheerin et al., 2021; Young et al., 2021; Qi et al., 2021; Hossack et al., 2020; Kuan et al., 2017). As a future work, we expect the proposed method can be extended to longitudinal data when such data become available.

In addition, the number of subject in the DNHS study is relatively small, and thus readers should be cautious when directly using the results that the DNHS dataset provides. In the future, with more date collected, we will apply the proposed method to a larger dataset. More discussion on the limitations of the DNHS data is provided in Section S.5.5 of Supplementary Materials.

## Acknowledgements

We would like to acknowledge support for this project from the National Institutes of Health grants R01MD011728, R01DA022720, and RC1MH088283; and from the National Science Foundation grants DMS-1821198 and DMS-1952406.

