## Supplementary Materials for "Heterogeneous Mediation Analysis on Epigenomic PTSD and Traumatic Stress in a Predominantly African American Cohort"

### S.1 Proofs

Note that when the distribution of $(X_i, \boldsymbol{Z}_i)$ is somewhat degenerate, different $\Theta$'s could result in the same distribution $f(X_i, \boldsymbol{M}_i, Y_i, \boldsymbol{Z}_i; \boldsymbol{\Theta}, C_i)$. For instance, if $X_i$ is constantly zero, then $f(X_i, \boldsymbol{M}_i, Y_i, \boldsymbol{Z}_i; \boldsymbol{\Theta}, C_i = h)$ does not depend on $\boldsymbol{b}_h$ or $\beta_h$ by equation (1). Similarly, if the first two components of $\boldsymbol{Z}_i$ are the same random variable, then changing $\boldsymbol{\Gamma}$ to a new matrix by swapping the first two columns does not affect $f(X_i, \boldsymbol{M}_i, Y_i, \boldsymbol{Z}_i; \boldsymbol{\Theta}, C_i)$. For this reason, we make the following non-degeneracy assumption.

**Condition 1** (Non-degeneracy)**.** *We suppose the distribution of* $(X_i, \boldsymbol{Z}_i)$ *is non-degenerate in the sense that* $f(X_i, \boldsymbol{M}_i, Y_i, \boldsymbol{Z}_i; \boldsymbol{\Theta}, C_i = h) = f(X_i, \boldsymbol{M}_i, Y_i, \boldsymbol{Z}_i; \boldsymbol{\Theta}', C_i = h)$ *implies* $(\boldsymbol{\Theta}_{1h}, \boldsymbol{\Theta}_2) = (\boldsymbol{\Theta}'_{1h}, \boldsymbol{\Theta}'_2)$.

*Proof of Proposition 1.* By the non-degeneracy condition, we have $\boldsymbol{\Theta}_2 = \boldsymbol{\Theta}'_2$ and $\boldsymbol{\Theta}_{1C_i} = \boldsymbol{\Theta}'_{1C'_i}$ for each $i$. It suffices to show that there is a permutation $\sigma$ on the set of group labels $\{1, 2, \cdots, H\}$ such that $\boldsymbol{\Theta}_{1h} = \boldsymbol{\Theta}'_{1\sigma(h)}$ for all $1 \leq h \leq H$.

For each $h$, there is some $i$ such that $C_i = h$ since each subgroup is nonempty. Define $\sigma(h) = C'_i$. We note that $\sigma$ is well-defined by showing that it does not depend on the choice of $i$. If $C_i = C_j = h$, then $\boldsymbol{\Theta}'_{1C'_j} = \boldsymbol{\Theta}_{1h} = \boldsymbol{\Theta}'_{1C'_i}$, and thus $C'_j = C'_i$.

The map $\sigma$ is injective for a similar reason as follows. Suppose $\sigma(h_1) = \sigma(h_2) = h'$. That is, there is some $i, j$ with $C_i = h_1$, $C_j = h_2$, and $C'_i = C'_j = h'$. Then we have $\boldsymbol{\Theta}_{1h_1} = \boldsymbol{\Theta}'_{1h'} = \boldsymbol{\Theta}_{1h_2}$ and thus $h_1 = h_2$ as desired.

As an injective self-map on a finite set, the map $\sigma$ is a permutation on the group labels, which has the desired property $\boldsymbol{\Theta}_{1h} = \boldsymbol{\Theta}'_{1\sigma(h)}$ by definition. This completes the proof. □

### S.2 Expanded Version of Algorithm 1

1. Set the number of subgroups $H$, tolerance $\epsilon$, and the tuning parameters.

2. Obtain initial subgroup labels $C_i^{(0)}$ for $i = 1, \ldots, N$, via the k-means clustering method (MacQueen, 1967) on all the variables.

   - Here the "all the variables" include the independent variable ($X_i$), pre-treatment confounders ($\boldsymbol{Z}_i$), potential mediators ($\boldsymbol{M}_i$), and the outcome variable ($Y_i$), where $i$ represents the $i$-th subject.

3. Calculate $\boldsymbol{\nu}^{(0)}$ based on $C_i^{(0)}$ for each subgroup separately.

   - The vector $\boldsymbol{\nu}$ consists of all the parameters in the following mediator and outcome models:

$$\boldsymbol{M}_i = \boldsymbol{b}_h X_i + \boldsymbol{\Gamma} \boldsymbol{Z}_i + \boldsymbol{\delta}_i, \tag{12}$$

$$Y_i = \beta_h X_i + \boldsymbol{\theta}_h^T \boldsymbol{M}_i + \boldsymbol{\gamma}^T \boldsymbol{Z}_i + \varepsilon_i, \tag{13}$$

   for each $h = 1, \ldots, H$, where $\boldsymbol{b}_h, \boldsymbol{\delta}_i, \boldsymbol{\theta}_h \in \mathbb{R}^p$, $\boldsymbol{\Gamma} \in \mathbb{R}^{p \times r}$ , and $\boldsymbol{\gamma} \in \mathbb{R}^r$ are parameters. Among these parameters, $\boldsymbol{b}_h$, $\beta_h$ and $\boldsymbol{\theta}_h$ are subgroup-specific, while the remaining ones are not. Based on initial subgroup labels $C_i^{(0)}$ for $i = 1, \ldots, N$, we can identify the initial subjects in each subgroup. To calculate the initial values of parameters $b_{h,j}$ (the $j$-th element in $\boldsymbol{b}_h$) and $\boldsymbol{\Gamma}_j$ (the $j$-th row of $\boldsymbol{\Gamma}$) for $1 \leq j \leq p$, we follow the mediator model in (12) by using the $j$-th mediator in $\boldsymbol{M}_i$ as a response variable, and the independent variable $X_i$ and pre-treatment confounders $\boldsymbol{Z}_i$ as covariates. With the initial subjects assigned in the $h$-th subgroup ($1 \leq h \leq H$), we calculate the ordinary least squares (OLS) estimator of $b_{h,j}$'s ($j = 1, \ldots, p$) as its initial value. If the number of initial subjects within the subgroup is smaller than the number of covariates, we use the Lasso estimator (Tibshirani, 1996) as its initial value. Since $\boldsymbol{\Gamma}$ is not subgroup-specific, we take an average of the estimators (OLS or Lasso) of $\boldsymbol{\Gamma}_j$ from all subgroups as the initial value.

Similarly, for the initial values of $\beta_h$, $\boldsymbol{\theta}_h$, and $\boldsymbol{\gamma}$, we follow the outcome model in (13) by using the outcome variable $Y_i$ as a response variable and $X_i$, $\boldsymbol{M}_i$, and $\boldsymbol{Z}_i$ as covariates. Based on subjects initially assigned to the $h$-th subgroup, we use the OLS estimators of $\beta_h$ and $\boldsymbol{\theta}_h$ as their initial values, or use the Lasso estimator if there is not a sufficient number of initial subjects within the subgroup. Since $\boldsymbol{\gamma}$ is not subgroup-specific, we use an average of the estimators of $\boldsymbol{\gamma}$ from all subgroups as its initial value.

4. At the $m$-th iteration, given $\boldsymbol{\nu}^{(m-1)}$ from the $(m-1)$-th iteration:

   (a) Calculate $\boldsymbol{\mu}^{(m-1)} \in \partial f_2(\boldsymbol{\nu}^{(m-1)})$.

   - Here $\boldsymbol{\mu}$ is defined as one element in the subdifferential of $f_2(\boldsymbol{\nu})$:
$$\partial f_2(\boldsymbol{\nu}) = \sum_{i=1}^{N} \text{co}\left\{\cup_{k \in J(\boldsymbol{\nu})} \partial F_{ik}(\boldsymbol{\nu})\right\} + N \sum_{h=1}^{H} \partial p_d(\boldsymbol{\Theta}_{1h}),$$
where $f_2(\boldsymbol{\nu})$ is defined in equation (9), $p_d(\boldsymbol{\Theta}_{1h}) = \beta_h^2/(a-1) + 2c_0^2\lambda_1(\|\boldsymbol{\theta}_h\|_2^2 + \|\boldsymbol{b}_h\|_2^2)$, $F_{ik}(\boldsymbol{\nu}) = \sum_{1 \leq h \leq H, h \neq k} L_{i,h}(\boldsymbol{\Theta}_{1h}, \boldsymbol{\Theta}_2)$, $J(\boldsymbol{\nu}) = \{1 \leq k \leq H : F_{ik}(\boldsymbol{\nu}) = \max_{1 \leq l \leq H} F_{il}(\boldsymbol{\nu})\}$, and "co" stands for a convex hull. In fact, since $p_d$ and $F_{ik}(\boldsymbol{\nu})$ are differentiable, the choice of $\boldsymbol{\mu}$ is not unique when there are multiple elements in the set $J(\boldsymbol{\nu})$; that is, when there are multiple subgroups $k$'s such that each $F_{ik}(\boldsymbol{\nu})$ can achieve the maximum $\max_{1 \leq l \leq H} F_{il}(\boldsymbol{\nu})$.
In our implementation, we use the first subgroup which can achieve this maximum. In fact, in practice, we have a unique choice for $\boldsymbol{\mu}$ according to $J(\boldsymbol{\nu})$ in most situations. We illustrate this with simulations in Section S.3.3.

(b) Solve $\boldsymbol{\nu}^{(m)}$ in (10) through the gradient descent.

- Recall that the parameter vector is $\boldsymbol{\nu} = (\beta_1, \boldsymbol{\theta}_1^T, \boldsymbol{b}_1^T, \dots, \beta_H, \boldsymbol{\theta}_H^T, \boldsymbol{b}_H^T, \boldsymbol{\gamma}^T, \boldsymbol{\Gamma}_1, \dots, \boldsymbol{\Gamma}_p)^T$, where $\boldsymbol{\Gamma}_j$ denotes the $j$-th row of $\boldsymbol{\Gamma}$ for $j = 1, \dots, p$. Explicitly, at the $m$-th iteration of the entire proposed algorithm, the objective function in the gradient descent algorithm is

$$
\begin{aligned}
& \tilde{f}^{(m)}(\boldsymbol{\nu}) \\
= \; & \sum_{i=1}^{N}\sum_{h=1}^{H}[\text{tr}\{(\boldsymbol{M}_i - \boldsymbol{b}_h X_i - \boldsymbol{\Gamma}\boldsymbol{Z}_i)(\boldsymbol{M}_i - \boldsymbol{b}_h X_i - \boldsymbol{\Gamma}\boldsymbol{Z}_i)^T\} + (Y_i - \beta_h X_i - \boldsymbol{\theta}_h^T \boldsymbol{M}_i \\
& -\boldsymbol{\gamma}^T \boldsymbol{Z}_i)^2] + N\sum_{h=1}^{H}\left\{\frac{\beta_h^2}{a-1} + 2c_0^2\lambda_1(\|\boldsymbol{\theta}_h\|_2^2 + \|\boldsymbol{b}_h\|_2^2) + p_{SCAD,\lambda_1,a}(\beta_h)\right. \\
& \left. +\lambda_1\sum_{j=1}^{p}\left(1 - \frac{1}{(1+c_0|b_{h,j}|)(1+c_0|\theta_{h,j}|)}\right)\right\} + \lambda_2 N\left\{\|\boldsymbol{\gamma}\|_1 + \sum_{j=1}^{p}\|\boldsymbol{\Gamma}_j\|_1\right\} \\
& +\lambda_0 N\left\{(\boldsymbol{\eta}^*)^T \boldsymbol{D}\boldsymbol{\nu} - \frac{\rho}{2}\|\boldsymbol{\eta}^*\|_2^2\right\} - f_2^{(m)}(\boldsymbol{\nu}),
\end{aligned}
$$

where $\boldsymbol{\eta}^* = S(\boldsymbol{D}\boldsymbol{\nu}/\rho)$, $\boldsymbol{D}$ is a difference operator corresponding to the pairwise differences on parameters in $\sum_{1\le h_1,h_2\le H}[|\beta_{h_1} - \beta_{h_2}| + \sum_{j=1}^{p}\{|\theta_{h_1,j} - \theta_{h_2,j}| + |b_{h_1,j} - b_{h_2,j}|\}]$,

$$
S(x) = \begin{cases} x, & -1 \le x \le 1, \\ 1, & x > 1, \\ -1, & x < -1, \end{cases}
$$

$$
p_{SCAD,\lambda_1,a}(\beta_h) = \begin{cases} \lambda_1|\beta_h|, & \text{if } |\beta_h| \le \lambda_1, \\ -\frac{|\beta_h|^2 - 2a\lambda_1|\beta_h| + \lambda_1^2}{2(a-1)}, & \text{if } \lambda_1 < |\beta_h| \le a\lambda_1, \\ \frac{(a+1)\lambda_1^2}{2}, & \text{if } |\beta_h| > a\lambda_1, \end{cases}
$$

and

$$
\begin{aligned}
& f_2^{(m)}(\boldsymbol{\nu}) \\
= \; & f_2(\boldsymbol{\nu}^{(m-1)}) + \left\langle \boldsymbol{\nu} - \boldsymbol{\nu}^{(m-1)}, \boldsymbol{\mu}^{(m-1)} \right\rangle \\
= \; & \sum_{i=1}^{N} \max_{1 \le k \le H} \left\{ \sum_{1 \le h \le H, h \ne k} \Big[ \mathrm{tr}\{ (\boldsymbol{M}_i - \boldsymbol{b}_h^{(m-1)} X_i - \boldsymbol{\Gamma}^{(m-1)} \boldsymbol{Z}_i)(\boldsymbol{M}_i - \boldsymbol{b}_h^{(m-1)} X_i \right. \\
& \left. - \boldsymbol{\Gamma}^{(m-1)} \boldsymbol{Z}_i)^T \} + (Y_i - \beta_h^{(m-1)} X_i - (\boldsymbol{\theta}_h^{(m-1)})^T \boldsymbol{M}_i - (\boldsymbol{\gamma}^{(m-1)})^T \boldsymbol{Z}_i)^2 \Big] \right\} \\
& + N \sum_{h=1}^{H} \left\{ \frac{(\beta_h^{(m-1)})^2}{a-1} + 2c_0^2 \lambda_1 (\|\boldsymbol{\theta}_h^{(m-1)}\|_2^2 + \|\boldsymbol{b}_h^{(m-1)}\|_2^2) \right\} \\
& + \left\langle \boldsymbol{\nu} - \boldsymbol{\nu}^{(m-1)}, \boldsymbol{\mu}^{(m-1)} \right\rangle .
\end{aligned}
$$

Here $\boldsymbol{\mu}^{(m-1)}$ is an element in the subdifferential of $f_2(\boldsymbol{\nu}^{(m-1)})$:

$$
\partial f_2(\boldsymbol{\nu}^{(m-1)}) = \sum_{i=1}^{N} \mathrm{co} \left\{ \cup_{k \in J(\boldsymbol{\nu}^{(m-1)})} \partial F_{ik}(\boldsymbol{\nu}^{(m-1)}) \right\} + N \sum_{h=1}^{H} \partial p_d(\boldsymbol{\Theta}_{1h}^{(m-1)}),
$$

where $p_d(\boldsymbol{\Theta}_{1h}) = \beta_h^2/(a-1) + 2c_0^2\lambda_1(\|\boldsymbol{\theta}_h\|_2^2 + \|\boldsymbol{b}_h\|_2^2)$, $J(\boldsymbol{\nu}) = \{1 \le k \le H : F_{ik}(\boldsymbol{\nu}) = \max_{1 \le l \le H} F_{il}(\boldsymbol{\nu})\}$, $F_{ik}(\boldsymbol{\nu}) = \sum_{1 \le h \le H, h \ne k} L_{i,h}(\boldsymbol{\Theta}_{1h}, \boldsymbol{\Theta}_2)$, and "co" stands for a convex hull. Specifically, note that

$$
\begin{aligned}
& \partial F_{ik}(\boldsymbol{\nu}) \\
= \; & -2 \left[ (Y_i - \beta_1 X_i - \boldsymbol{\theta}_1^T \boldsymbol{M}_i - \boldsymbol{\gamma}^T \boldsymbol{Z}_i)(X_i, \boldsymbol{M}_i^T), (\boldsymbol{M}_i - \boldsymbol{b}_1 X_i - \boldsymbol{\Gamma} \boldsymbol{Z}_i)^T X_i, \ldots, \right. \\
& (Y_i - \beta_H X_i - \boldsymbol{\theta}_H^T \boldsymbol{M}_i - \boldsymbol{\gamma}^T \boldsymbol{Z}_i)(X_i, \boldsymbol{M}_i^T), (\boldsymbol{M}_i - \boldsymbol{b}_H X_i - \boldsymbol{\Gamma} \boldsymbol{Z}_i)^T X_i, \\
& \sum_{1 \le h \le H, h \ne k} \left\{ (Y_i - \beta_h X_i - \boldsymbol{\theta}_h^T \boldsymbol{M}_i - \boldsymbol{\gamma}^T \boldsymbol{Z}_i) \boldsymbol{Z}_i^T, (M_{i1} - b_{h,1} X_i - \boldsymbol{\Gamma}_1 \boldsymbol{Z}_i) \boldsymbol{Z}_i^T, \ldots, \right. \\
& \left. \left. (M_{ip} - b_{h,p} X_i - \boldsymbol{\Gamma}_p \boldsymbol{Z}_i) \boldsymbol{Z}_i^T \right\} \right]^T
\end{aligned}
$$

is a $\{(1+2p)H + r + pr\}$-dimensional vector, and

$$
\sum_{h=1}^{H} \partial p_d(\boldsymbol{\Theta}_{1h}) = 2[\beta_1/(a-1), 2c_0^2\lambda_1(\boldsymbol{\theta}_1^T, \boldsymbol{b}_1^T), \ldots, \beta_H/(a-1), 2c_0^2\lambda_1(\boldsymbol{\theta}_H^T, \boldsymbol{b}_H^T), \mathbf{0}_{r+pr}]^T
$$

is also a $\{(1+2p)H + r + pr\}$-dimensional vector, where $M_{ij}$ and $b_{h,j}$ are the $j$-th elements in the vectors $\boldsymbol{M}_i$ and $\boldsymbol{b}_h$, respectively, for $j = 1, \ldots, p$; and $\mathbf{0}_{r+pr}$ is a $(r+pr)$-dimensional zero vector.

- At the $m$-th iteration, the gradient of the objective function $\tilde{f}^{(m)}(\boldsymbol{\nu})$ is

$$\nabla \tilde{f}^{(m)}(\boldsymbol{\nu}) = g_1(\boldsymbol{\nu}) + N g_2(\boldsymbol{\nu}) + \lambda_0 N \boldsymbol{D}^T \boldsymbol{\eta}^* - \boldsymbol{\mu}^{(m-1)}, \tag{14}$$

where

$$\begin{aligned} & g_1(\boldsymbol{\nu}) \\ = \; & -2\sum_{i=1}^{N} \Big[(Y_i - \beta_1 X_i - \boldsymbol{\theta}_1^T \boldsymbol{M}_i - \boldsymbol{\gamma}^T \boldsymbol{Z}_i)(X_i, \boldsymbol{M}_i^T), (\boldsymbol{M}_i - \boldsymbol{b}_1 X_i - \boldsymbol{\Gamma} \boldsymbol{Z}_i)^T X_i, \ldots, \\ & (Y_i - \beta_H X_i - \boldsymbol{\theta}_H^T \boldsymbol{M}_i - \boldsymbol{\gamma}^T \boldsymbol{Z}_i)(X_i, \boldsymbol{M}_i^T), (\boldsymbol{M}_i - \boldsymbol{b}_H X_i - \boldsymbol{\Gamma} \boldsymbol{Z}_i)^T X_i, \\ & \sum_{h=1}^{H} \big\{(Y_i - \beta_h X_i - \boldsymbol{\theta}_h^T \boldsymbol{M}_i - \boldsymbol{\gamma}^T \boldsymbol{Z}_i)\boldsymbol{Z}_i^T, (M_{i1} - b_{h,1} X_i - \boldsymbol{\Gamma}_1 \boldsymbol{Z}_i)\boldsymbol{Z}_i^T, \ldots, \\ & (M_{ip} - b_{h,p} X_i - \boldsymbol{\Gamma}_p \boldsymbol{Z}_i)\boldsymbol{Z}_i^T\big\}\Big]^T, \end{aligned}$$

$$\begin{aligned} & g_2(\boldsymbol{\nu}) \\ = \; & \Bigg[\frac{2\beta_1}{a-1} + \lambda_1 \left\{ I(\beta_1 \le \lambda_1) + \frac{(a\lambda_1 - \beta_1)_+}{(a-1)\lambda_1} I(\beta_1 > \lambda_1) \right\}, \\ & 4c_0^2 \lambda_1 \theta_{1,1} + \frac{\lambda_1 c_0 \operatorname{sign}(\theta_{1,1})}{(1 + c_0|b_{1,1}|)(1 + c_0|\theta_{1,1}|)^2}, \ldots, \\ & 4c_0^2 \lambda_1 \theta_{1,p} + \frac{\lambda_1 c_0 \operatorname{sign}(\theta_{1,p})}{(1 + c_0|b_{1,p}|)(1 + c_0|\theta_{1,p}|)^2}, \\ & 4c_0^2 \lambda_1 b_{1,1} + \frac{\lambda_1 c_0 \operatorname{sign}(b_{1,1})}{(1 + c_0|b_{1,1}|)^2(1 + c_0|\theta_{1,1}|)}, \ldots, \\ & 4c_0^2 \lambda_1 b_{1,p} + \frac{\lambda_1 c_0 \operatorname{sign}(b_{1,p})}{(1 + c_0|b_{1,p}|)^2(1 + c_0|\theta_{1,p}|)}, \\ & \ldots, \\ & \frac{2\beta_H}{a-1} + \lambda_1 \left\{ I(\beta_H \le \lambda_1) + \frac{(a\lambda_1 - \beta_H)_+}{(a-1)\lambda_1} I(\beta_H > \lambda_1) \right\}, \\ & 4c_0^2 \lambda_1 \theta_{H,1} + \frac{\lambda_1 c_0 \operatorname{sign}(\theta_{H,1})}{(1 + c_0|b_{H,1}|)(1 + c_0|\theta_{H,1}|)^2}, \ldots, \\ & 4c_0^2 \lambda_1 \theta_{H,p} + \frac{\lambda_1 c_0 \operatorname{sign}(\theta_{H,p})}{(1 + c_0|b_{H,p}|)(1 + c_0|\theta_{H,p}|)^2}, \\ & 4c_0^2 \lambda_1 b_{H,1} + \frac{\lambda_1 c_0 \operatorname{sign}(b_{H,1})}{(1 + c_0|b_{H,1}|)^2(1 + c_0|\theta_{H,1}|)}, \ldots, \\ & 4c_0^2 \lambda_1 b_{H,p} + \frac{\lambda_1 c_0 \operatorname{sign}(b_{H,p})}{(1 + c_0|b_{H,p}|)^2(1 + c_0|\theta_{H,p}|)}, \\ & \lambda_2 \left\{\operatorname{sign}(\boldsymbol{\gamma}^T), \operatorname{sign}(\boldsymbol{\Gamma}_1), \ldots, \operatorname{sign}(\boldsymbol{\Gamma}_p)\right\}\Bigg]^T, \end{aligned}$$

and $\text{sign}(\cdot)$ takes the signs of a vector component-wisely. The third term on the right-hand side of equation (14) follows from Chen et al. (2012, Theorem 1).

- At the $l$-th iteration of the gradient descent algorithm, we update $\boldsymbol{\nu}_{(l)} = \boldsymbol{\nu}_{(l-1)} - t_l \nabla \tilde{f}^{(m)}(\boldsymbol{\nu}_{(l-1)})$ until convergence, where the step size $t_l$ is determined by the following backtracking line search (Stanimirović and Miladinović, 2010):
  i. Set numbers $t_0 > 0$, $0 < c_1 < 0.5$ and $c_1 < c_2 < 1$.
  ii. Let $t = t_0$
  iii. While $\tilde{f}^{(m)}(\boldsymbol{\nu}_{(l-1)}) - \tilde{f}^{(m)}(\boldsymbol{\nu}_{(l-1)} - t\nabla \tilde{f}^{(m)}(\boldsymbol{\nu}_{(l-1)})) < c_1 t \|\nabla \tilde{f}^{(m)}(\boldsymbol{\nu}_{(l-1)})\|_2^2$, we let $t = c_2 t$. Otherwise, stop.
  iv. Return $t_l = t$.

(c) Update subgroup labels $C_i^{(m)}$ based on the loss function in (3) for $i = 1, \ldots, N$.

- We calculate the loss function of each observation with parameters over different subgroups, and then assign the observation or subject to the subgroup which minimizes the loss for that observation or subject.
- This subgroup identification is not done through post-hoc clustering. Instead, the subgroup labels and other parameters are both estimated through minimizing the objective function in the DC iterations. Specifically, at the $m$-th iteration, the subgroup label $C_i^{(m)}$ of the $i$-th subject is estimated as the subgroup which gives the minimum loss for the subject based on the updated parameters $\boldsymbol{\nu}^{(m)}$ in Step 4b. Thus, the subgroup identification fits into the DC algorithm.

5. Iterate Step 4 until $\|\boldsymbol{\nu}^{(m)} - \boldsymbol{\nu}^{(m-1)}\|_1 < \epsilon$

### S.3 Additional Simulations

#### S.3.1 Larger-scale Setting

We conduct a simulation setting with larger sample size $N$ and larger number of potential mediators $p$. Specifically, in the following Setting 4, we simulate data with $N = 500$ and $p = 700$.

**Setting 4.** *We investigate a larger-scale high-dimensional case proceeding similarly as in Setting 3 except that* $N = 500$*,* $n_1 = 200$*,* $n_2 = 300$*, and* $p = 700$*.*

The results of Setting 4 are in Table 7, showing that the proposed method produces much smaller overall false rates (FN+FP) for mediator selection and smaller MSE for heterogeneous mediation effects than the HIMA. Thus the proposed algorithm can be applied to datesets with larger $N$ and $p$ to facilitate the detection of heterogeneous sub-population effects. In addition, the results of the proposed method are much better than those in Setting 3 possibly due to higher sparsity under Setting 4, since we increase the sample size and the number of potential mediators but do not increase the number of true mediators in Setting 4.

Moreover, we observe that the computational time of the proposed algorithm is within one hour in a personal laptop for one replication in Setting 4, including tuning all hyperparameters. The proposed method converges much faster in other settings since the corresponding dimensions $N$ and $p$ are much smaller. For example, the computational time is within 20 minutes for one replication including all the parameter tuning in Setting 3. Therefore, the proposed method can be expected to converge in a reasonable time.

#### S.3.2 Different Initial Values

Since the proposed objective function in (6) is non-convex, we acknowledge that the optimization problem depends on starting values. However, as shown in Tables 2–4, with the "warm-up" initial values in Algorithm 1, the estimator of the proposed method performs much better than existing estimators in most cases, indicating that the proposed estimator is a "good" solution.

We also acknowledge that, as a non-convex optimization, with different initial values, the proposed algorithm might not always converge to a unique value. However, the algorithm is still stable to some extent. We conduct simulations with different initial values under Settings 1 and 2. Specifically, in Setting 1 (homogeneous structure), we obtain two solutions with the same initial clusters but different initial values for parameters $\boldsymbol{b}_1$ and $\boldsymbol{\theta}_1$: One solution has the true values of the parameters as initial values, and the other one has the true values plus one as initial values. Note

Table 7: FN, FP, FN+FP, and MSE under Setting 4. "HIMA" stands for the high-dimensional mediation analysis method. The "$\theta_{1s}$" and "$\theta_{2s}$" represent the signal strength in $\boldsymbol{\theta}_1$ and $\boldsymbol{\theta}_2$, respectively, and "$\rho$" is a correlation parameter.

| | $(\theta_{1s}, \theta_{2s})$ | Method | FN | FP | FN+FP | MSE |
|---|---|---|---|---|---|---|
| $\rho = 0$ | (0.5, -0.5) | **Proposed** | 0.002 | 0.000 | 0.002 | 0.000 |
| | | HIMA | 0.853 | 0.001 | 0.854 | 0.004 |
| | (0.8, -0.8) | **Proposed** | 0.001 | 0.000 | 0.001 | 0.001 |
| | | HIMA | 0.945 | 0.001 | 0.946 | 0.009 |
| | (1, -1) | **Proposed** | 0.003 | 0.000 | 0.003 | 0.001 |
| | | HIMA | 0.969 | 0.000 | 0.969 | 0.014 |
| | (4, -4) | **Proposed** | 0.004 | 0.001 | 0.005 | 0.018 |
| | | HIMA | 0.994 | 0.000 | 0.994 | 0.228 |
| $\rho = 0.2$ | (0.5, -0.5) | **Proposed** | 0.009 | 0.000 | 0.009 | 0.001 |
| | | HIMA | 0.896 | 0.001 | 0.897 | 0.004 |
| | (0.8, -0.8) | **Proposed** | 0.005 | 0.000 | 0.005 | 0.001 |
| | | HIMA | 0.987 | 0.000 | 0.987 | 0.009 |
| | (1, -1) | **Proposed** | 0.011 | 0.000 | 0.011 | 0.001 |
| | | HIMA | 0.993 | 0.000 | 0.993 | 0.014 |
| | (4, -4) | **Proposed** | 0.011 | 0.000 | 0.011 | 0.021 |
| | | HIMA | 0.982 | 0.000 | 0.982 | 0.228 |

that the true values of parameters are sparse in Setting 1. Thus, these two sets of initial values are quite different, as the former one is sparse, while the latter one is dense.

We calculate the average rate of equality of the two solutions in terms of parameters $\boldsymbol{b}_1$ and $\boldsymbol{\theta}_1$ across $100$ replications. The rates are provided in Table 8. We can see that the rates are all greater than $82\%$, implying that the solutions with two different initial values are the same in most situations. Moreover, for the heterogeneous Setting 2, we obtain two solutions with the same initial values for parameters but different initial clusters. Similarly, we check the equality of the two solutions and find they are the same in all $100$ replications.

#### S.3.3 Non-uniqueness of $\mu$

As mentioned in Section S.2, the choice of $\boldsymbol{\mu}$ is not unique when there are multiple elements in the set $J(\boldsymbol{\nu})$. To investigate the non-uniqueness of $\boldsymbol{\mu}$, we conduct simulations under Setting 2 of Section 4. With each signal strength, we calculated the set $J(\boldsymbol{\nu})$ more than $320,000$ times on

Table 8: The rate of equality of two solutions with different initial values under Setting 1 based on 100 replications. The "$\theta_{1s}$" represents the signal strength in $\boldsymbol{\theta}_1$.

| $\theta_{1s}$ | $\boldsymbol{b}_1$ | $\boldsymbol{\theta}_1$ |
|---|---|---|
| 0.2 | 0.980 | 0.929 |
| 0.3 | 1.000 | 0.910 |
| 0.4 | 0.980 | 0.828 |

Table 9: FN, FP, FN+FP, and MSE of the proposed method with another choice for $\boldsymbol{\mu}$ under Setting 2. The "$\theta_{1s}$" and "$\theta_{2s}$" represent the signal strengths in $\boldsymbol{\theta}_1$ and $\boldsymbol{\theta}_2$, respectively.

| $(\theta_{1s}, \theta_{2s})$ | FN | FP | FN+FP | MSE |
|---|---|---|---|---|
| (0.5, -0.5) | 0.034 | 0.007 | 0.041 | 0.004 |
| (1, -1) | 0.003 | 0.001 | 0.004 | 0.006 |
| (4, -4) | 0.001 | 0.000 | 0.001 | 0.061 |

average per replication. Among these, the proportion of $J(\boldsymbol{\nu})$ containing multiple elements is less than $1\%$. That is, the non-uniqueness of $\boldsymbol{\mu}$ occurs rarely in practice.

In addition, we implement the proposed method with another choice for $\boldsymbol{\mu}$ under Setting 2. Specifically, we use the last subgroup instead of the first subgroup among all the groups which achieve the maximum of $F_{ik}(\boldsymbol{\nu})$ when calculating $\boldsymbol{\mu}$. The results are provided in Table 9, which are similar to the original results of the proposed method with the first subgroup, as shown in Table 3. Thus, the implementation results are quite robust to the choice of $\boldsymbol{\mu}$.

#### S.3.4 Various Coefficients

We conduct simulations with various coefficients in the following settings. The results are provided in Table 10, which show that the proposed method still produces much smaller FN+FP and MSE than the existing HIMA method when the coefficients are widely different.

**Setting 5.** *We generate data similarly as in Setting 1 except* $(\theta_{1,1}, \theta_{1,2}, \theta_{1,3}, \theta_{1,4}) = (0.2, 0.4, 0.6, 1.2)$.

**Setting 6.** *We generate data similarly as in Setting 2 except* $(\theta_{1,1}, \theta_{1,2}, \theta_{1,3}) = (0.5, 1, 1.5)$ *and* $(\theta_{2,1}, \theta_{2,2}, \theta_{2,3}) = (-0.5, -1, -1.5)$.

**Setting 7.** *We generate data similarly as in Setting 3 except* $(\theta_{1,1}, \theta_{1,2}, \theta_{1,3}) = (0.5, 1, 1.5)$ *and* $(\theta_{2,1}, \theta_{2,2}, \theta_{2,3}) = (-0.5, -1, -1.5)$.

Table 10: The FN, FP, FN+FP, and MSE under Settings 5-7, where "HIMA" stands for the high-dimensional mediation analysis method.

| Settings | Method | FN | FP | FN+FP | MSE |
|---|---|---|---|---|---|
| Setting 5 | **Proposed** | 0.003 | 0.000 | 0.003 | 0.002 |
| | HIMA | 0.178 | 0.000 | 0.178 | 0.002 |
| Setting 6 | **Proposed** | 0.035 | 0.000 | 0.035 | 0.010 |
| | HIMA | 0.813 | 0.003 | 0.816 | 0.163 |
| Setting 7, $\rho = 0$ | **Proposed** | 0.397 | 0.001 | 0.398 | 0.011 |
| | HIMA | 0.953 | 0.001 | 0.954 | 0.036 |
| Setting 7, $\rho = 0.2$ | **Proposed** | 0.376 | 0.000 | 0.376 | 0.010 |
| | HIMA | 0.980 | 0.000 | 0.980 | 0.035 |

#### S.3.5 Moderation

We consider simulations where the underlying conceptual model involves a moderator. Specifically, we let the underlying model be a moderation model and a moderated mediation model in the following Settings 8 and 9, respectively. The corresponding results are provided in Table 11.

In Setting 8, since there is just a moderation model for $Y_i$, the potential mediators $\boldsymbol{M}_i$ are independent of the outcome and the independent variable, implying that there is no true mediator. Then we do not need to calculate any false negative (FN) rates. As shown in Table 11, the false positive (FP) rates and the mean-squared-errors (MSE) of the proposed method and the HIMA method are similar and all close to 0. Thus, the proposed method and the HIMA method both perform well under the moderation model.

As for Setting 9, the underlying model is a moderated mediation one. Then there are several true mediators. The results in Table 11 show that the proposed method performs much better than the HIMA method in terms of FN+FP and MSE.

**Setting 8.** *We generate data similarly as in Setting 2 in Section 4, except that* $\boldsymbol{M}_i = \boldsymbol{\delta}_i$ *and* $Y_i = 2X_i + 2X_iR_i + R_i + \varepsilon_i$ *for* $i = 1, \ldots, N$*, where* $R_i$ *is a moderator taking value* $0$ *for* $50$

*subjects and* $1$ *for remaining subjects.*

**Setting 9.** *We generate data similarly as in Setting 2 except that* $\boldsymbol{M}_i = \boldsymbol{b}X_iR_i + \boldsymbol{\delta}_i, Y_i = \beta X_iR_i + \boldsymbol{\theta}^T\boldsymbol{M}_iR_i + \varepsilon_i$*, where* $R_i$ *is a moderator taking value* $1$ *for* $50$ *subjects, and* $-1$ *for the remaining subjects. We let* $\boldsymbol{b} = (1, 1, 1, 0, \ldots, 0)^T$*,* $\boldsymbol{\theta} = (\theta_s, \theta_s, \theta_s, 0, \ldots, 0)^T$*,* $\beta = 0.5$*, where* $\theta_s = 0.5, 1,$ *or* $4$*.*

Table 11: The FN, FP, FN+FP, and MSE under Settings 8 and 9, where "HIMA" stands for the high-dimensional mediation analysis method.

| Settings | Method | FN | FP | FN+FP | MSE |
|---|---|---|---|---|---|
| Setting 8 | **Proposed** | — | 0.002 | 0.002 | 0.000 |
| | HIMA | — | 0.001 | 0.001 | 0.000 |
| Setting 9, $\theta_s = 0.5$ | **Proposed** | 0.057 | 0.001 | 0.058 | 0.005 |
| | HIMA | 0.983 | 0.001 | 0.984 | 0.025 |
| Setting 9, $\theta_s = 1$ | **Proposed** | 0.006 | 0.000 | 0.006 | 0.006 |
| | HIMA | 0.960 | 0.000 | 0.960 | 0.099 |
| Setting 9, $\theta_s = 4$ | **Proposed** | 0.000 | 0.000 | 0.000 | 0.058 |
| | HIMA | 0.773 | 0.000 | 0.773 | 1.509 |

#### S.3.6 Non-normality of the Independent Variable

To investigate the potential impact of non-normality of the independent variable, we conduct simulations under the following Setting 10, where we generate the independent variable from a skewed Laplace distribution. The results are provided in Table 12. Compared to the performance of the proposed method with normally distributed variables (see results in Table 3), the proposed method performs slightly worse in the non-normality case. However, the proposed method still performs much better than the existing HIMA method in terms of both FN+FP and MSE here. Therefore, the proposed method is robust against the normality of the independent variable.

**Setting 10.** *We generate data similarly as in Setting 2 except that we generate* $X_i$ *from an asymmetric Laplace distribution with the location parameter* $0.5$*, scale parameter* $0.2$*, and the skewness parameter* $0.7$*. for* $i = 1, \ldots, N$*.*

Table 12: FN, FP, FN+FP, and MSE under Setting 10. "HIMA" stands for the high-dimensional mediation analysis method. The "$\theta_{1s}$" and "$\theta_{2s}$" represent the signal strength in $\boldsymbol{\theta}_1$ and $\boldsymbol{\theta}_2$, respectively.

| $(\theta_{1s}, \theta_{2s})$ | Method | FN | FP | FN+FP | MSE |
|---|---|---|---|---|---|
| (0.5, -0.5) | **Proposed** | 0.140 | 0.007 | 0.147 | 0.007 |
| | HIMA | 0.960 | 0.003 | 0.963 | 0.025 |
| (1, -1) | **Proposed** | 0.026 | 0.002 | 0.028 | 0.012 |
| | HIMA | 0.743 | 0.003 | 0.746 | 0.093 |
| (4, -4) | **Proposed** | 0.006 | 0.000 | 0.006 | 0.131 |
| | HIMA | 0.667 | 0.001 | 0.668 | 1.456 |

### S.4 Additional Real Data Analyses

#### S.4.1 Common Patterns in Subgroup Identification

Among the 100 random splits, the proposed method identifies three subgroups in 58 replications, two subgroups in 3 replications, and one subgroup in 39 replications. Thus, in most split training datasets, the proposed method identifies three subgroups, which is to some extent consistent with the number of subgroups identified by the proposed method based on the whole dataset.

Moreover, for the subgroups based on the whole dataset, 573 pairs of within-group subjects are also clustered to the same subgroup in all 58 replications where three subgroups are identified, while this does not occur for any cross-group pair of subjects. In addition, 1422 cross-group pairs in the whole-dataset analysis are clustered to different subgroups in all these 58 replications, while this only occurs for 3 within-group pairs. Therefore, there are common patterns in the analyses of the 100 random split datasets and the whole dataset, which essentially indicates the stability of the proposed method.

#### S.4.2 Racial Make-up

The racial make-up of the three identified subgroups based on the entire dataset is provided in Table 13, which shows that the subjects are not grouped based on race since each subgroup contains both AAs and European Americans. There is no significant difference on the distribution of race across these three subgroups, indicating that the heterogeneity in mediation is not due to race. Thus we include all subjects instead of AAs alone for more powerful analysis.

Table 13: Racial make-up of subgroups. “AA” represents African American. “EA” represents European American.

| | AA | EA | Other |
|---|---|---|---|
| Subgroup 1 | 51 | 8 | 1 |
| Subgroup 2 | 24 | 2 | 0 |
| Subgroup 3 | 36 | 3 | 0 |

### S.5 Additional Discussion

#### S.5.1 Subgroup Identification

Our proposed subgroup identification is different from that in the individualized-multi-directional method (IMDM) of Tang et al. (2021). Compared with the IMDM in Tang et al. (2021), the proposed method has two main advantages. First, the proposed method is more parsimonious with a simpler subgroup structure and thus can be more stable. The IMDM allows different subgroup partitions with different heterogeneous-effect predictors, and there is no borrowed information across subgrouping on different predictors in the IMDM. In contrast, the proposed method focuses on subject-based subgrouping, and pursues one-dimensional subgrouping only. Second, the proposed method does not require that homogeneous predictors be pre-specified, while the IMDM does. In the proposed method, the homogeneous predictors are identified in a data-driven way via a fused lasso penalty.

The disadvantage of the proposed method is that the subgrouping in the proposed method is less flexible than that in the method of Tang et al. (2021). With only one-dimensional subgrouping on all predictors across subjects, it might be difficult for the proposed method to handle situations where the subgrouping varies greatly across predictors.

Since we consider a high-dimensional mediation framework which has already involved many parameters even without heterogeneity, we adopt the proposed method instead of that in Tang et al.

(2021) to focus on subject-wise subgrouping.

#### S.5.2 Exposure–Mediator Interaction

We did not allow any exposure–mediator interaction in our current study, which is often accounted for in causal mediation (Valeri and VanderWeele, 2013; Liu et al., 2016; Rijnhart et al., 2021). In fact, it is challenging to add interaction terms since we have already had high-dimensional potential mediators. On the other hand, to capture data heterogeneity, many existing studies propose to add interaction terms into models (Shuster and van Eys, 1983; Gail and Simon, 1985; Gunter et al., 2011). These, however, rely on certain prior knowledge and additional model assumptions, such as a linear relationship between the response and some pre-specified interactions. Yet, it could be impractical to impose which interaction terms should be included in the model prior to analysis, while simply adding all interaction terms of all predictors would apparently yield an ultra-high dimensionality of unknown parameters in the model and thus likely suffers from over-fitting. In addition, the heterogeneity of mediation effects could be due to more complex reasons, which might not be captured by the interaction terms. In contrast, the proposed method detects heterogeneous mediation effects without imposing additional assumptions.

#### S.5.3 Latent Subpopulations

The proposed heterogeneous-effect model is inspired by relevant background knowledge and the reasonable scientific hypothesis that the potential mediator effects are not necessarily the same for the entire population. Some scientific work has provided evidence that mediation effects could be different corresponding to different groups of individuals (Qin and Hong, 2017; Dyachenko and Allenby, 2018). For example, Dyachenko and Allenby (2018) found that all subjects in a real case study can be separated into a mediating group and a non-mediating group, and the mediating variable only has a non-zero effect in the mediating group.

Therefore, in this paper, we model the possible heterogeneous mediation effects by leveraging the subpopulation structure. One key advantage of the proposed method is that the latent subgroups are not pre-specified but rather identified completely based on the data. Thus, our method does not rely on any prior knowledge on subpopulations.

In fact, we do not require the existence of heterogeneity or multiple subpopulations. Our model just provides a way to consider multiple subpopulations if heterogeneity exists. The homogeneous model is a special case of our model with only one subpopulation. In the implementation of our model, the number of subgroups is selected in a data-driven fashion. Thus, we are able to identify the homogeneous setting when there is only one subgroup.

For example, Setting 1 in the simulation studies (Section 4) generates a homogeneous underlying true model with only one sub-population. To investigate whether the proposed method can identify the homogeneity, we calculate the correct rates of selecting subgroup number under Setting 1, which is the proportion of replications where the number of subgroups is correctly selected via the proposed method. The results are provided in Table 5, where $\theta_{1s}$ represents the magnitude of mediation effects. Our numerical results show that the proposed method correctly determines the number of subgroups in most situations, especially when the mediation effects are relatively large.

For the situation with relatively small mediation effects ($\theta_{1s} = 0.2$), even if the correct rate of selecting the subgroup number does not achieve $100\%$, the false negative rates (FN) and the false positive rates (FP) for mediator selection are less than $0.05$, and the FN+FP is much smaller than that of the existing high-dimensional mediation analysis (HIMA) method. The corresponding mean-squared-errors (MSE) of mediation effects are also small. These results are provided in Table 2.

Lastly, we acknowledge that the assessment of identified subpopulations is very challenging and still remains an open issue. In recent years, some efforts have been made to address post-subgrouping inference issues. For example, Guo and He (2021) proposed a de-biasing bootstrap inference procedure to test the selected subgroups with heterogeneous treatment effects, and Wang et al. (2021) suggested the application of that procedure in mediation analysis for testing the validity of selected subgroups. In addition, some downstream analysis can be further conducted to investigate the potential factors that are relevant to subgrouping, such as exploring the demographical variables and/or other factors across different subgroups. This will also provide further implications for prediction on future observations.

#### S.5.4 Moderated Mediation and Mediated Moderation

There are many psychology publications on moderated mediation and mediated moderation (Muller et al., 2005; Morgan-Lopez and MacKinnon, 2006; Edwards and Lambert, 2007; Bucy and Tao, 2007). In the following, we first introduce the definitions of mediated moderation and moderated mediation. Suppose that $Y$ is an outcome variable, $X$ is an independent variable, $M$ is a mediator variable, and $R$ is a moderator variable. According to Muller et al. (2005), the underlying models for both mediated moderation and moderated mediation are

$$Y = \alpha_{10} + \alpha_{11}X + \alpha_{12}R + \alpha_{13}XR + \varepsilon_{01}, \tag{15}$$

$$M = \alpha_{20} + \alpha_{21}X + \alpha_{22}R + \alpha_{23}XR + \varepsilon_{02}, \tag{16}$$

$$Y = \alpha_{30} + \alpha_{31}X + \alpha_{32}R + \alpha_{33}XR + \alpha_{34}M + \alpha_{35}MR + \varepsilon_{03}. \tag{17}$$

The model in (15) allows the overall effect of the independent variable $X$ on the outcome $Y$ to be moderated by the moderator $R$. The model in (16) allows the effect of the independent variable $X$ on the mediator $M$ to be moderated, and the model in (17) allows both the effects of the mediator and the independent variable on the outcome to be moderated.

Mediated moderation is applicable when overall moderation occurs (i.e., $\alpha_{13} \neq 0$) and a mediator accounts for the moderation/interaction $XR$ (i.e., $\alpha_{33}$ is smaller in absolute value than $\alpha_{13}$). Moderated mediation occurs when the mediation effect varies across levels of the moderator variable $R$. Specifically, the effect of independent variable $X$ on the mediator $M$ is moderated by $R$ in equation (16), and the effect of mediator $M$ on $Y$ is moderated in equation (17). The difference between the mediated moderation and moderated mediation is whether there is an overall moderation of the independent variable $X$ on the outcome $Y$ in the process. If there is, the process is mediated moderation; otherwise, it is moderated mediation (Muller et al., 2005).

In fact, mediated moderation and moderated mediation can be considered as a special case of heterogeneous mediation, since the mediation effect varies across subjects in mediated moderation and moderated mediation. The proposed heterogeneous mediation is more general and flexible as it does not require any pre-knowledge of moderators or interaction terms in the models, while mediated moderation and moderated mediation do require that.

#### S.5.5 DNHS Data Limitations

In the DNHS data, we collected the experienced traumas, the blood specimens for DNAm, and the PTSD Checklist (PCL) score of each subject nearly at the same time, where the PCL score measures the recent symptoms of the subject. It is very likely that the traumas occur before the collection of DNAm and PCL scores. Since the methylation levels are stable in a short term (Byun et al., 2012; Zaimi et al., 2018), it is also likely that the change of methylation due to traumas occurs before PTSD symptoms. Nevertheless, information about the temporal order is lacking in this DNHS study.

Moreover, we acknowledge that the sample size of our real data application is modest. In fact, many existing studies on PTSD-related mediation analysis have about one hundred participants (Kwon et al., 2021; Ruhlmann et al., 2019; Demir et al., 2020; Kearney et al., 2013; van der Vleugel et al., 2020; Kelly et al., 2019). A relatively small sample size could be a challenge for existing statistical methods, which motivates us to develop more powerful statistical methods to identify mediators with heterogeneity and high-dimensional issues. As shown in our simulation studies (e.g., Setting 3) in Section 4, the proposed method outperforms existing methods in terms of both selection of mediators and estimation of mediation effects, when the sample size ($N = 100$) and the number of potential mediators ($p = 150$) are similar to those in the real data application ($N = 125, p = 144$).

Also, in our current study, we only have records of trauma types for experienced traumas without any other categories such as "heard about" or "witnessed" traumas. Our future work with larger datasets would be better positioned to incorporate more nuanced analyses of trauma types into the model.
